# Genomic hypomethylation in cell-free DNA predicts responses to checkpoint blockade in lung and breast cancer

**DOI:** 10.1101/2023.10.31.565052

**Authors:** Kyeonghui Kim, Hyemin Kim, Inkyung Shin, Seung-Jae Noh, Jeong Yeon Kim, Koung Jin Suh, Yoo-Na Kim, Jung-Yun Lee, Dae-Yeon Cho, Se Hyun Kim, Jee Hyun Kim, Se-Hoon Lee, Jung Kyoon Choi

## Abstract

Genomic hypomethylation has recently been identified as a determinant of therapeutic responses to immune checkpoint blockade (ICB). However, tumor tissue is often unattainable, and tissue-based methylation profiling suffers from low tumor purity. In this study, we developed an assay named iMethyl to estimate the genomic hypomethylation status from cell-free DNA (cfDNA) as well as tissue by deep targeted sequencing of young LINE-1 elements with > 400,000 reads per sample. iMethyl was applied to a total of 653 ICB samples encompassing lung cancer (cfDNA n=167; tissue n=137; cfDNA early during treatment n=40), breast cancer (cfDNA n=91; tissue n=50; PBMC n=50; cfDNA at progression n=44), and ovarian cancer (tissue n=74). iMethyl-tissue had better predictive power than tumor mutation burden and PD-L1 expression. Furthermore, iMethyl-liquid predicted ICB responses accurately regardless of the tumor purity of tissue samples. iMethyl-liquid was also able to monitor therapeutic responses early during treatment (3 or 6 weeks after initiation of ICB) and detect progressive hypomethylation accompanying tumor progression. In conclusion, our method allows for reliable noninvasive prediction, early evaluation, and monitoring of clinical responses to ICB therapy.

## Introduction

Immune checkpoint blockade (ICB) therapy has proven to be effective in multiple cancer types and is widely being used clinically. However, only a subset of patients receiving ICB therapy experience a durable clinical benefit. There is therefore an imperative need for biomarkers that can predict its therapeutic responses. Various biomarkers have been proposed as determinants of treatment efficacy including tumor mutation burden (TMB) (1–3), expression of inhibitory targets such as PD-L1 (4–6), and defects in particular pathways (7), but with limited accuracy and clinical utility.

Cell-free DNA (cfDNA) refers to extracellular fragments of DNA found in plasma and body fluid. With advances in sequencing technology, capturing genetic or epigenetic features from plasma cfDNA at a high resolution is emerging as an alternative to tissue biopsies. cfDNA is easily obtainable whereas tissue sampling often requires invasive measures such as surgery. Moreover, cfDNA may be unaffected by confounding factors such as tumor purity that obfuscate the interpretation of tumor biopsies. Efforts are being made to apply cfDNA in cancer immunotherapy; cfDNA load (8–11), TMB (11–13), and copy number instability (14–16) have been identified as markers for the monitoring or prediction of clinical responses to ICB treatment. However, no attempts have been made to use cfDNA methylation in cancer immunotherapy.

In our previous work (17), we have shown that genomic methylation loss, especially in late-replicating domains (18), is coupled with immune evasion of tumors due to the silencing of genes involved in antigen processing and presentation, major histocompatibility complex, and cytokine-cytokine receptor interaction by inducing promoter hypermethylation. Therefore, genomic hypomethylation estimated by array probes mapping to evolutionarily young subfamilies of LINE-1 elements (L1HS and L1PA) was able to predict the clinical benefit of ICB therapy more accurately than TMB in multiple lung cancer and melanoma cohorts (17).

In this work, we hypothesized that LINE-1 methylation levels could be captured from cfDNA for the prediction of ICB responses. Based on this assumption, we developed an assay named iMethyl (immune-Methyl) based on high-depth targeted sequencing of the genomic regions corresponding to the 10 LINE-1 array probes that had been validated in our previous work (17). To test its utility for noninvasive prognosis in immunotherapy, we applied iMethyl to a total of 653 ICB samples encompassing lung cancer (cfDNA n=167; tissue n=137; cfDNA early during treatment n=40), breast cancer (cfDNA n=91; tissue n=50; PBMC n=50; cfDNA at progression n=44), and ovarian cancer (tissue n=74). In particular, we sought to find the advantage of applying iMethyl to cfDNA (iMethyl-liquid) over applying it to tissue specimen (iMethyl-tissue). We also tested iMethyl-liquid as a tool for monitoring disease progression during ICB therapy. Our results collectively suggest that iMethyl-liquid is a robust and accurate method for noninvasive prediction and monitoring of clinical responses to ICB treatment.

## Results

### Development of iMethyl and comparison with array data

Progressive methylation loss occurs due to the failure of methylation maintenance machinery to remethylate newly synthesized stands during DNA replication (19). This genomic demethylation process is especially observed in late-replicating partial methylation domains (PMDs) of rapidly dividing cancer genomes (18). To quantitatively capture this process, we measured the CpG methylation levels of the open sea probes across the whole genome and those that lie within PMDs from array data for tissue samples in lung cancer. This whole-genome or PMD-based measurement of genomic hypomethylation was well represented by the average methylation level of the young LINE-1 probes used in our previous work (17) (Supplementary Figure 1A). Importantly, these LINE-1 probes could differentiate clinical responses to ICB therapy better than the whole-genome or PMD-based measurements (Supplementary Figure 1B).

Here, we sought to develop an assay based on the targeted sequencing of the loci corresponding to these array probes (Supplementary Tables 1-2). The probe sequences were estimated to map approximately 2,000 locations across the human genome (17). Thus, to achieve a 0.5% resolution of methylation measurement across a broad range of these different LINE-1 copies, we aimed at > 400,000 reads over all target loci combined for each sample.

To validate its usage as a standalone assay using our lung cancer ICB cohort (Supplementary Table 3), iMethyl was first performed for tissue samples (n=137, Supplementary Table 4) and compared with the array LINE-1 data (n=60, Supplementary Table 5). Unlike the array platform, our sequencing-based assay could capture cytosine methylation signals from both CpG and CpA sites for the three non-CpG probes: P8, P9, and P10 (Figure 1A and Supplementary Figure 2). The principal component analysis highlighted the differences between the iMethyl and array data mainly caused by the three probes and P5 (Figure 1B). These differences resulted in different probe-wise clustering of the matched samples (n=36, Supplementary Figure 3).

**Figure 1.**
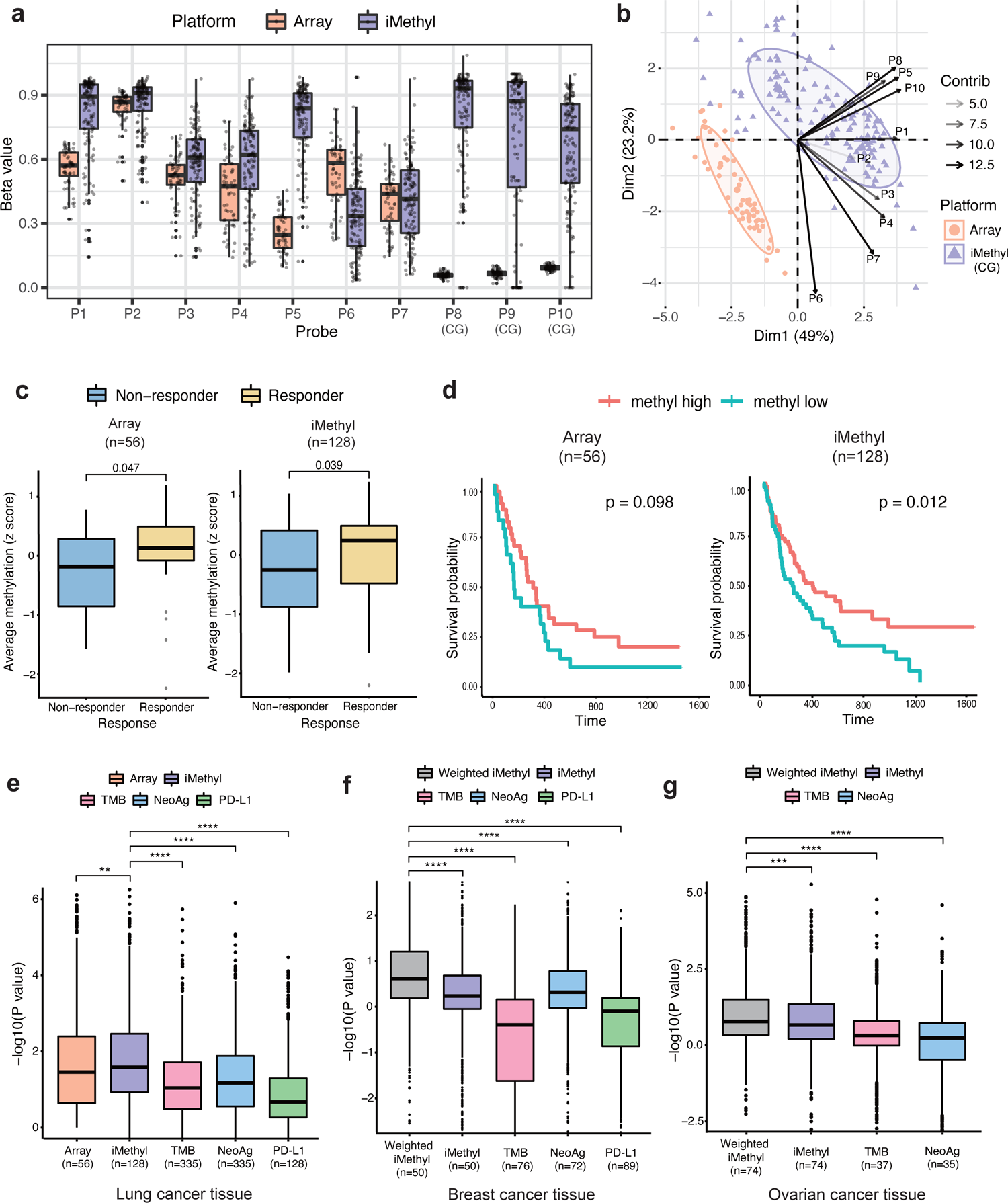
Comparison of iMethyl with array and other ICB markers in tissue samples. (A) Distribution of beta values per probe by array versus iMethyl in the tissue samples of our lung cancer cohort. For iMethyl on non-CpG probes (P8, P9, and P10), methylation signals from only CpG sequences are plotted. (B) Principal component analysis of the beta values of the array and iMethyl probes. (C) Comparison of the average methylation values of the array and iMethyl probes between the non-responders and responders of our lung cancer cohort. (D) Survival analysis between the methylation-high and -low group based on the average methylation values of the array and iMethyl probes in our lung cancer cohort. (E-G) Prediction power for the clinical outcome of ICB therapy comparing iMethyl, TMB, neoantigen load (NeoAg), and PD-L1 expression in our (E) lung cancer, (F) breast cancer, and (G) ovarian cancer cohort. For each cohort, the performance of the predictors was estimated by 1,000 times of bootstrapping of individual patient samples. In each sampling, the P value from the survival analysis was obtained. The 1,000 P values for the different predictors were compared by the Wilcoxon signed-rank test (P value of **** < 1×10^-4^, *** < 1×10^-3^, ** < 1×10^-2^).

### iMethyl-tissue outperforms other measures in predicting ICB responses

We investigated iMethyl’s ability to discriminate therapeutic responses to ICB after excluding samples without clinical information. The average methylation level of the 10 probes was lower in the non-responders of our lung cancer cohort (Figure 1C), supporting the contribution of genomic methylation loss to the immune evasion of tumors (17). Our survival analysis also illustrates that samples with low methylation levels have a worse prognosis (Figure 1D). Importantly, the iMethyl measures outperformed the array-based readouts of the LINE-1 probes in both analyses (Figure 1C-D).

We then sought to validate iMethyl as a predictive marker for ICB therapy by using our breast cancer cohort (Supplementary Table 6) and ovarian cancer cohort (Supplementary Table 7). iMethyl was performed for breast cancer tissues (n=50, Supplementary Table 8) and ovarian cancer tissues (n=74, Supplementary Table 9). Whole-exome sequencing and immunohistochemistry were also performed to obtain previous predictive measures such as TMB, neoantigen load (NeoAg), and PD-L1 expression.

For a more robust statistical analysis, we implemented a bootstrapping technique by resampling patients in each cohort 1,000 times, performing the survival analysis for each resample, and comparing the distribution of the P values from the 1,000 survival analyses. According to the distribution of the P values, iMethyl-tissue outperformed TMB, NeoAg, and PD-L1 expression in discriminating clinical responses across all cohorts (Figure 1E-G). Taken together, at the tissue level, the 10 iMethyl probes could adequately represent genomic methylation loss as well as serve as an accurate biomarker for predicting ICB responses.

### iMethyl-liquid uncouples the confounding effects of tumor purity from tissue methylomes

To test whether iMethyl-liquid can represent tumor tissue methylomes, we performed iMethyl with cfDNA (n=167, Supplementary Table 10) samples in our lung cancer cohort (Supplementary Table 3), and with cfDNA (n=91, Supplementary Table 11), PBMC (n=50, Supplementary Table 12), and tissue (n=50, Supplementary Table 8) samples in our breast cancer cohort (Supplementary Table 6). The LINE-1 methylation values from the different sources showed distinctive distributions, with cfDNA ranging higher than tissue and lower than PBMC (Figure 2A). Whereas iMethyl-PBMC indicated invariably high LINE-1 methylation in most samples, iMethyl-liquid represented a similar degree of intertumoral diversity as iMethyl-tissue (Figure 2A-B). In contrast to iMethyl-PBMC, iMethyl-liquid showed significant correlations with iMethyl-tissue (Supplementary Figure 4), resulting in similar clustering patterns (Supplementary Figure 5) across the matched samples.

**Figure 2.**
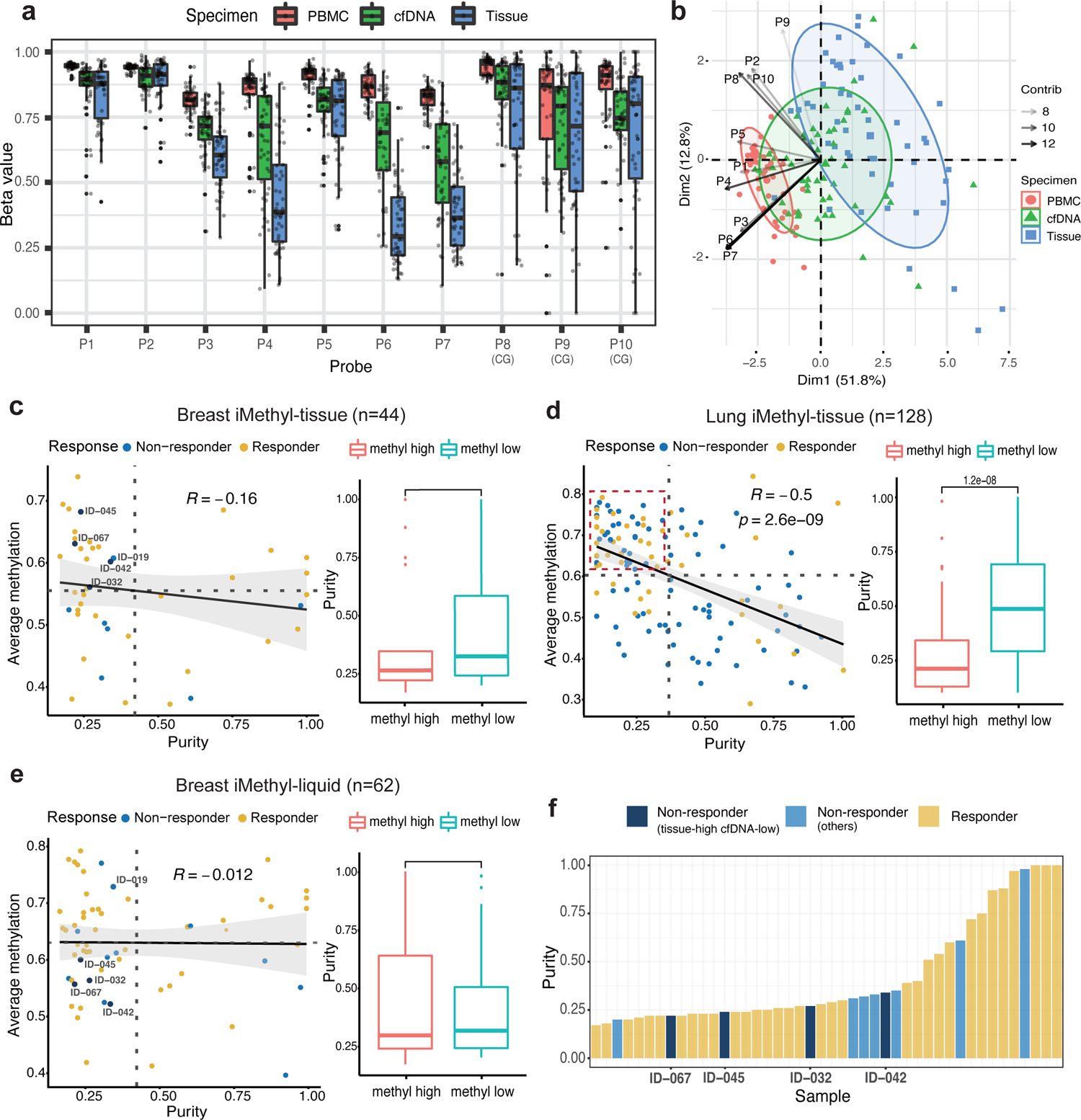
Evaluation of iMethyl-liquid using matched tissue and PBMC samples. (A) Distribution of beta values per probe by matched iMethyl-tissue, -liquid, and -PBMC for our breast cancer samples. (B) Principal component analysis of the beta values of iMethyl from three specimen sources. (C-D) Influence of tumor purity on the average methylation level from iMethyl-tissue in our (C) breast cancer cohort and (D) lung cancer cohort examined by the correlation analysis (left) and by comparing the estimate of tumor purity between the methylation-high and -low group (right). ICB non-responders with high iMethyl-tissue and low tumor purity are marked (C) with their patient ID or (D) by the red box. (E) Influence of tumor purity on the average methylation level from iMethyl-liquid in our breast cancer cohort. ICB non-responders with high iMethyl-tissue and low tumor purity are marked with their patient ID. (F) Tumor purity of the four tissue-high and cfDNA-low non-responder samples in comparison with the remaining samples of our breast cancer cohort.

Tumor purity is a critical issue in cancer genomics and epigenomics (20,21). Cellular heterogeneity of a tissue specimen can often blur tumor characteristics. cfDNA, on the other hand, can provide a more unbiased estimation in proportion to tumor burden. To investigate the confounding effects of tumor purity, we utilized the whole-exome sequencing data of our lung and breast cancer cohorts. Expectedly, the tissue methylation levels were overestimated in samples with low tumor purity in both cancer types (Figure 2C-D); there were multiple cases of ICB non-responders that would be misclassified by iMethyl-tissue because of its overestimation (blue dots marked with their patient ID or enclosed by the red box). In sharp contrast, iMethyl-liquid was not affected by tumor purity (Figure 2E). Indeed, four of the five non-responders with high iMethyl-tissue and low purity had iMethyl-liquid lower than the cohort average (Figure 2E) and also than expected by iMethyl-tissue (Supplementary Figure 6). Therefore, for certain samples with low purity, resistance to ICB therapy coupled with cancer-specific genomic methylation loss can only be predicted by cfDNA analysis (Figure 2F).

As illustrated by these samples, tissue methylation may have limited accuracy compared with cfDNA methylation unless tumor purity is completely accounted for. Indeed, in breast cancer, iMethyl-liquid showed statistical power in stratifying prognoses (P=0.0017) after excluding one sample without clinical information whereas iMethyl-tissue failed to yield reliable clinical prediction (P=0.48) (Figure 3A left). Our bootstrap analysis confirmed the superior performance of iMethyl-liquid over iMethyl-tissue (Figure 3A right). The same patterns were recapitulated when using only the matched samples (Supplementary Figure 7) and also with the lung cancer data (Figure 3B). Taken together, iMethyl-liquid yields more accurate interpretation of tumor methylomes and thus a more accurate prediction of ICB responses, thanks to its independence of tumor purity.

**Figure 3.**
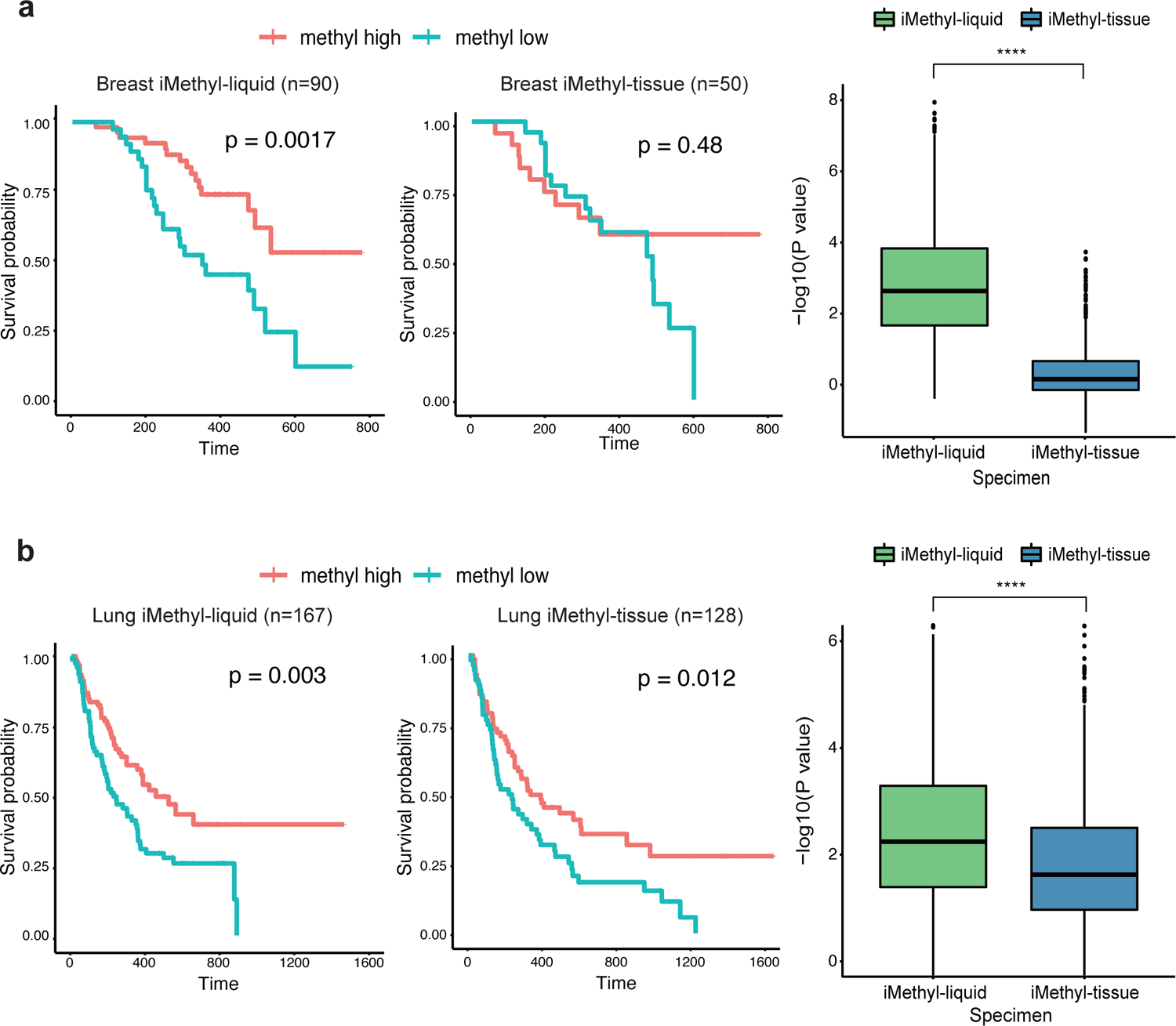
Comparison of iMethyl-liquid and iMethyl-tissue in predicting ICB efficacy. (A-B) Survival analysis (left) and bootstrap analysis (right) using our (A) breast cancer and (B) lung cancer cohort samples. The survival analysis was performed for the methyl-high and - low group to compare iMethyl-liquid and iMethyl-tissue. For the bootstrap analysis, performance was estimated by 1,000 trials of resampling of individual patient samples. In each sampling, the P value from the survival analysis was obtained. The resulting 1,000 P values for iMethyl-liquid and iMethyl-tissue were compared by the Wilcoxon signed-rank test (P value of **** < 1×10^-4^).

### Implementation of probe weighting improves iMethyl performance

To improve the performance of iMethyl as an ICB efficacy predictor, we tested applying the weighted average of the iMethyl probes. For iMethyl-tissue, we trained probe weights for optimization of predicting lung cancer ICB responses and applied the optimal weights to the 10 probe signals of breast cancer and ovarian cancer samples. Although the weight training was performed using completely independent samples, this process improved predictive power compared to averaging without weighting (gray versus violet in Figure 1F-G). The improvements made by applying the weights across cancer types were also evident in survival analyses (Supplementary Figure 8). iMethyl-tissue was also improved by training probe weights on iMethyl-liquid of the same cancer type in both ICB cohorts (Supplementary Figure 9).

Finally, we concentrated on how the performance of iMethyl-liquid can be improved by probe weighting. Compared to no weighting (Figure 4A), iMethyl-liquid weighted by iMethyl-liquid of a different cancer type (Figure 4B) or by iMethyl-tissue of the same cancer type (Figure 4C) showed increased performance in discriminating the better survival of high methylation tumors in response to ICB. In both ICB cohorts, our bootstrapping analyses involving 5,000 trials of resampling also showed improvements by cross-weighting across cancer types or specimen sources as well as by self-weighting (Figure 4D-E).

**Figure 4.**
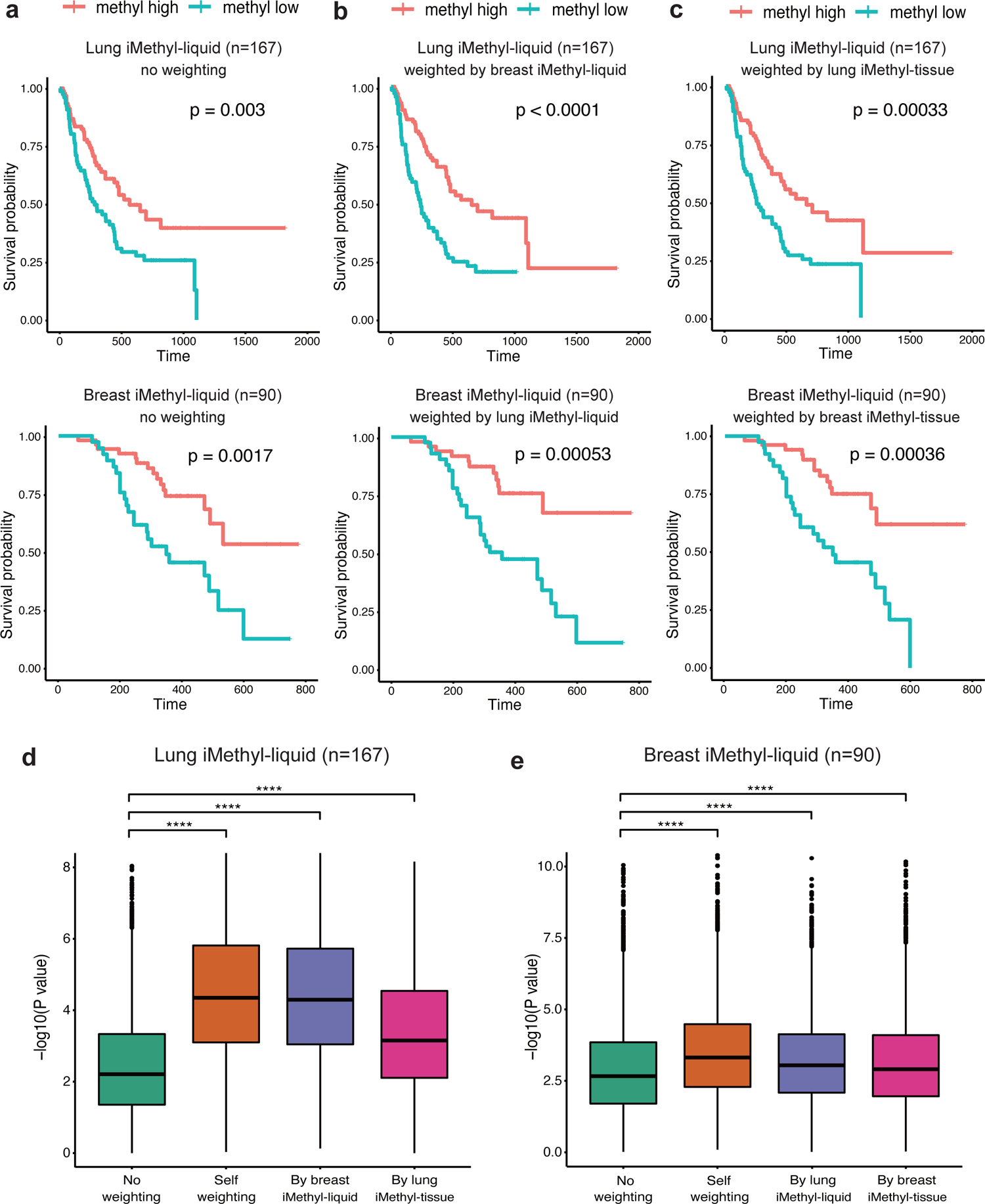
Enhanced prediction power of iMethyl-liquid by probe weighting. (A-C) Survival analysis between the methylation-high and -low group based on the average methylation values from iMethyl-liquid of our lung cancer samples (upper) and breast cancer samples (lower) with (A) no weighting, (B) weighting by iMethyl-liquid of a different cancer type, and (C) weighting by iMethyl-tissue of the same cancer type. (D-E) Prediction power for the clinical outcome of ICB therapy comparing different weighting schemes for (D) lung cancer iMethyl-liquid and (E) breast cancer iMethyl-liquid. Performance was estimated by 5,000 trials of bootstrapping of individual patient samples. In each sampling, the P value from the survival analysis was obtained. The resulting 5,000 P values for different weighting schemes were compared by the Wilcoxon signed-rank test (P value of **** < 1×10^-4^).

In addition, we compared the distribution of the actual methylation values with or without weighting between ICB responders and non-responders. Significantly lower methylation was observed for non-responders without any weighting (Supplementary Figure 10A); however, the methylation values weighted by iMethyl-liquid of a different cancer type (Supplementary Figure 10B) or those weighted by iMethyl-tissue of the same cancer type (Supplementary Figure 10C) showed a larger discrepancy between responders and non-responders in both ICB cohorts.

These results are significant because the weighting was trained on independent samples. Not only the weighting across cancer types (Figure 4B) but also across sample sources in lung cancer (Figure 4C upper) was based on different samples. Although cfDNA samples were weighted by a subset of matching tissue samples in our breast cancer cohort (Supplementary Table 6), iMethyl-liquid and iMethyl-tissue were performed for different samples of our lung cancer cohort (Supplementary Table 3). This implies that iMethyl-liquid can be further empowered by this cross-weighting approach while maintaining robustness to overfitting. For example, iMethyl-tissue data from pre-existing tissues of the same cohort can be utilized to refine the prediction of new samples by iMethyl-liquid.

### Monitoring of disease progression during ICB therapy by iMethyl-liquid

Another merit of cfDNA analysis is its noninvasive monitoring of disease progression. To illustrate the benefits from this capability, we used patient samples with progressive disease (n=44) in our breast cancer cohort (Supplementary Table 6). For these samples, iMethyl-liquid was performed at the point of disease progression (Supplementary Table 13) for comparison with matched iMethyl-liquid at the baseline (Supplementary Table 11). A differential methylation analysis of the matched samples showed significant methylation loss at most of the LINE-1 probes, especially P2, P3, P5, P8, and P9, but except for P4, P6, and P10, during tumor progression (Figure 5A). P2 and P8 showed the largest number of samples with a decrease in methylation and the lowest number of samples with an increase in methylation (Figure 5B). The average methylation level of P2, P3, P5, P8, and P9, that is, the probes with the most significant changes (Figure 5A), decreased overall at disease progression in most of the samples (Figure 5C).

**Figure 5.**
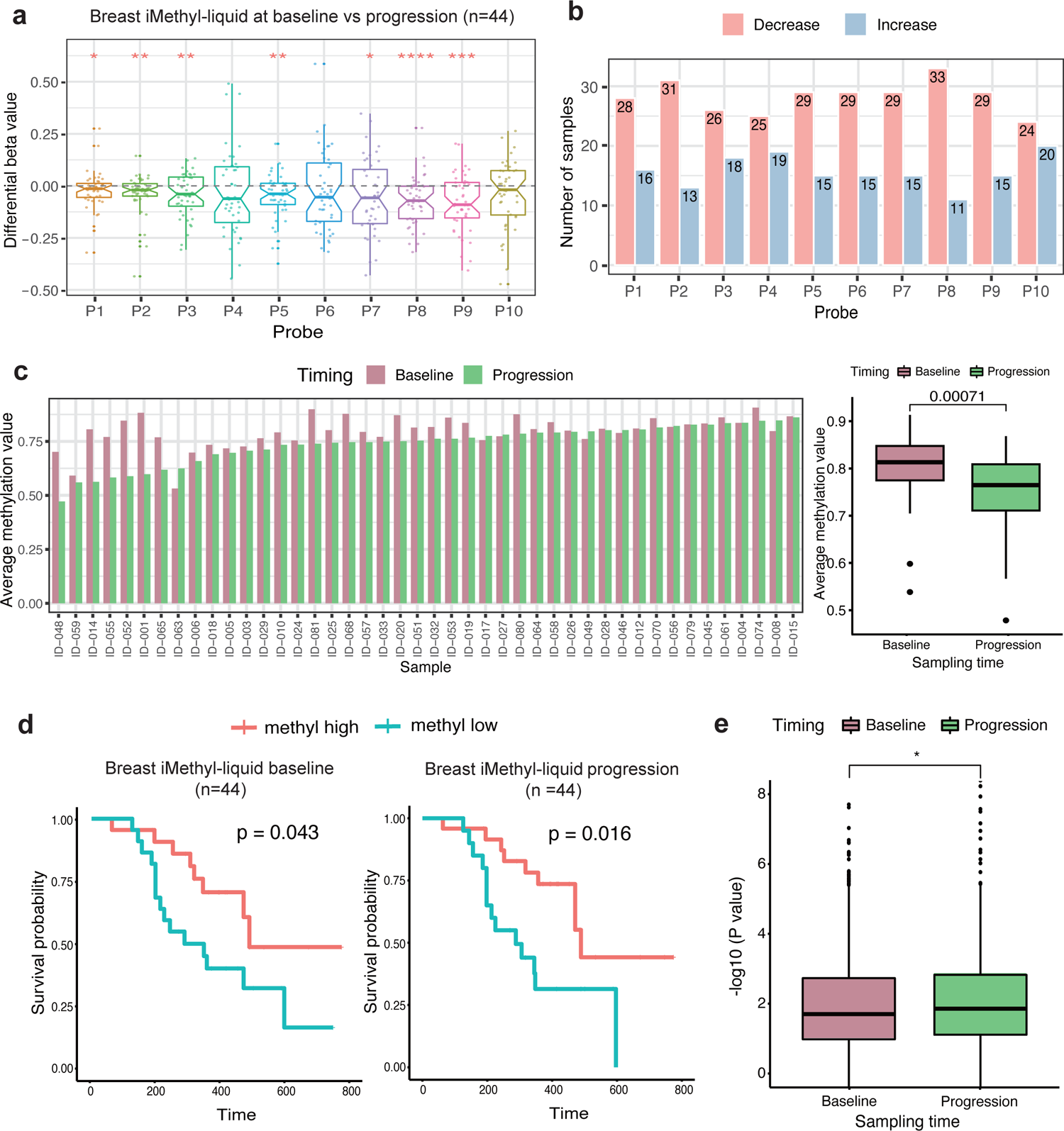
Comparison of baseline and post-progression iMethyl-liquid. (A) Differential beta values between matched baseline and progression iMethyl-liquid for our breast cancer samples (n=44). The differential beta values were tested to determine if they were less than zero by the one-sample t-test (P value of **** < 1×10^-4^, *** < 1×10^-3^, ** < 1×10^-2^, * < 5×10^-2^). (B) Number of samples whose methylation level at each probe increased or decreased at the point of tumor progression. (C) Average methylation value of the 5 selected probes, namely, P2, P3, P5, P8, and P9, compared between the baseline and progression point per sample (left) and for all samples (right). (D) Survival analysis between the methylation-high and -low group based on the average methylation values of all 10 probes from iMethyl-liquid at the baseline (left) and progression (right) point. (E) Prediction power for the clinical outcome of ICB therapy comparing the baseline and progression iMethyl-liquid. Performance was estimated by 1,000 trials of bootstrapping of individual patient samples. In each sampling, the P value from the survival analysis was obtained. The resulting 1,000 P values for the baseline and progression iMethyl-liquid were compared by the Wilcoxon signed-rank test (P=0.021).

When progression of the disease occurs, a clinical decision is often required regarding whether to continue the ongoing therapy or consider another therapeutic option. Therefore, we tested whether iMethyl-liquid retains its capability of predicting the ultimate outcome of ICB treatment at the point of disease progression. When matched with the survival data, the predictive power of iMethyl-liquid based on the average of the 10 probes was displayed at the progression point (P=0.016) as well as at the baseline (P=0.043) (Figure 5D). Our bootstrap analysis confirmed a slightly better performance of iMethyl-liquid at tumor progression than at the baseline (Figure 5E). Although P2, P3, P5, P8, and P9 showed the most significant changes during tumor progression, the iMethyl-liquid estimate based on the average of these 5 selected probes lost predictive power as compared to that based on the average of all the 10 probes (Figure 5D versus Supplementary Figure 11A). Our bootstrap analysis confirmed that using the average of all the probes predicts patient survival better (Supplementary Figure 11B). Although the 5 probes may reflect changes in tumor burden, the features of genomic hypomethylation associated with immune evasion and immunotherapeutic resistance may be better captured by the absolute methylation levels of all probes combined together.

These results demonstrate the utility of iMethyl-liquid in noninvasive monitoring of ICB responses based on the detection of progressive hypomethylation especially based on particular LINE-1 probes. Moreover, this approach may help with clinical decision-making according to the genomic hypomethylation status of samples at the point of disease progression.

### Early evaluation of ICB responses by iMethyl-liquid

As iMethyl-liquid showed its capability to monitor changes in the status of genomic methylation accompanying tumor progression, we next asked whether it could be also used to evaluate therapeutic responses early during treatment (EDT). To this end, we selected 20 responders and 20 non-responders from our lung cancer cohort (Supplementary Table 3) and generated iMethyl-liquid data for these samples at 3 weeks and 6 weeks after commencing treatment (Supplementary Table 14). In particular, we selected the samples such that the baseline (pretreatment) iMethyl-liquid readouts were similar between the responders and non-responders.

As done with the breast iMethyl-liquid data (Figure 5A), the differential methylation levels between pretreatment and EDT were computed for each probe (Supplementary Figure 12). The per-probe methylation changes compared between the responders and non-responders indicated an overall correlation between the favorable ICB response and the increase of the iMethyl-liquid level (Supplementary Figure 12). Notably, this correlation was more pronounced at the probes that showed significant methylation changes during breast cancer progression, except for P5 (Supplementary Figure 12 and Figure 6A). It remains to be tested whether the contradictory results for P5 differential methylation can be attributed to the tumor type difference (i.e., breast cancer versus lung cancer). In any case, the average methylation level of the remaining 4 probes, namely, P2, P3, P8, and P9, showed progressive hypomethylation in the non-responder samples whereas progressive hypermethylation in the responder samples (Figure 6A-6B).

**Figure 6.**
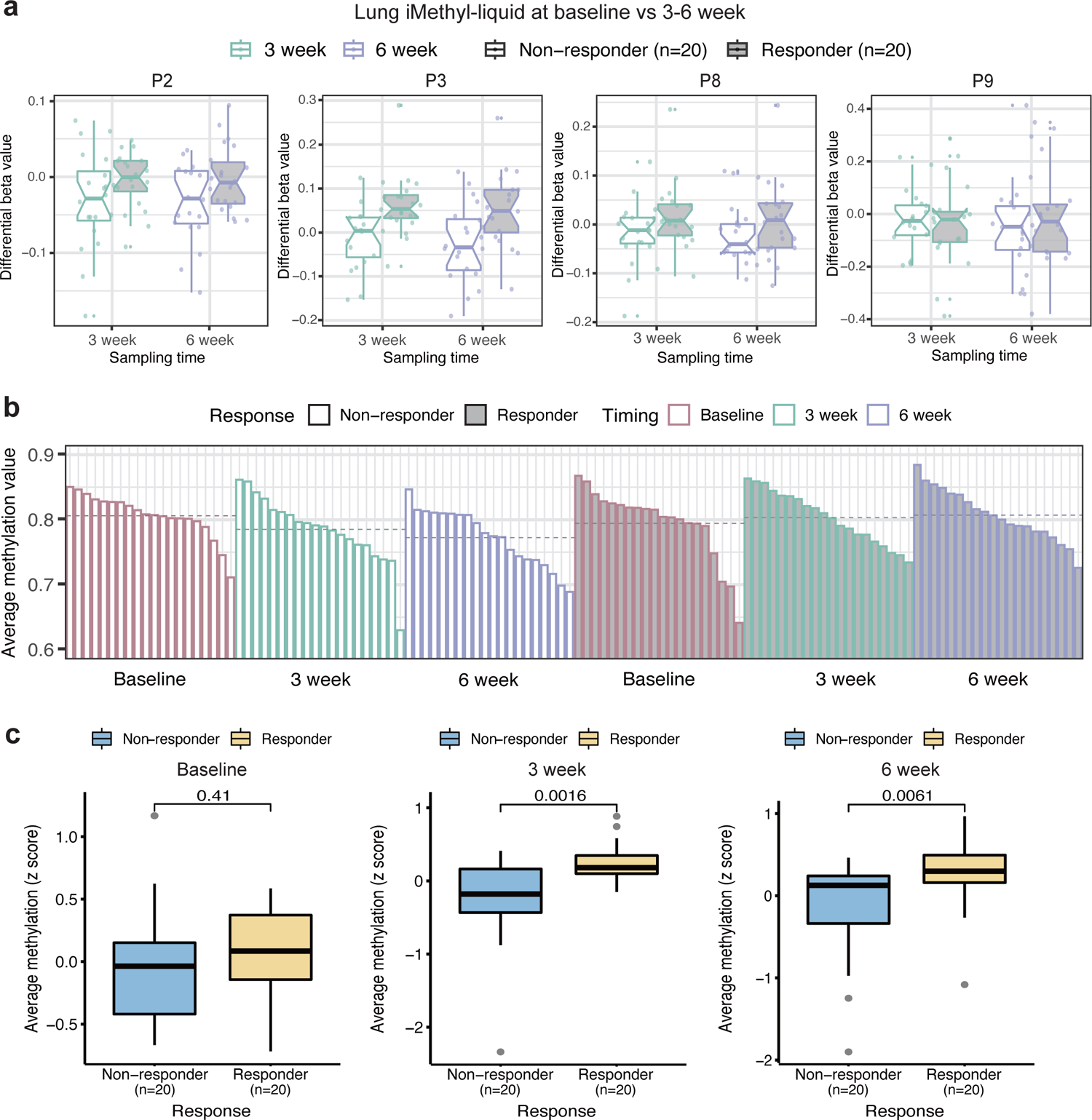
Differential iMethyl-liquid between pretreatment and EDT in association with clinical responses. (A) Differential beta values between matched pretreatment and EDT iMethyl-liquid in our lung cancer samples (20 responders and 20 non-responders) at P2, P3, P8, and P9 probes for pretreatment versus 3 weeks after initiating treatment (left) and for pretreatment versus 6 weeks after initiating treatment (right). (B) Average methylation value of the 4 probes compared between pretreatment and EDT per sample for the non-responders (left) and responder samples (right). Samples were ordered according to the methylation level within each time group. Horizontal dotted lines indicate the mean of the methylation levels for each time group. (C) Average methylation value of all 10 probes compared between the responders and non-responders at the baseline point (left), 3 weeks after initiation of ICB (middle), and 6 weeks after initiation of ICB (right).

As discussed in the previous section, genomic hypomethylation estimated by the average of all the probes may serve as a better predictor of ICB responses although differential methylation at only a subset of the probes seems to reflect changes in tumor burden. Therefore, we measured the average iMethyl-liquid signal of all the 10 probes at the baseline and at 3 or 6 weeks after the initiation of ICB treatment in order to evaluate its predictive power. The baseline measures were similar between the responding and non-responding groups because that was how the samples were selected (Figure 6C left). In contrast, the differences between the responders and non-responders were significantly pronounced in the EDT samples (Figure 6C middle and right).

These results illustrate the feasibility of measuring differential iMethyl-liquid for a noninvasive early evaluation of therapeutic responses to ICB. In addition, thanks to this feature of iMethyl-liquid reflecting tumor burden changes, a more accurate prediction of the ultimate clinical outcome may be made early during treatment than by using pretreatment samples alone.

## Discussion

In our previous work (17), we showed that genomic methylation aberration is an important marker of resistance to antitumor immunity in treatment-naïve samples as well as ICB-treated tumors. On the basis of large-scale TCGA and multiple ICB cohort data, we showed that rapidly dividing cells escape antitumor immune responses in association with genomic demethylation coupled with the silencing of critical genes involved in the response of tumors to host immune activity. Our data was based on genomic hypomethylation estimated by signals from array probes mapped to young subfamilies of LINE-1 elements (L1HS and L1PA) (17).

In the present work, we made improvements by first employing amplicon sequencing for genomic loci corresponding to these LINE-1 probes. When applied to the tumor tissues of our ICB cohorts, the assay named iMethyl resulted in 486,000 ∼ 930,000 sequencing reads per sample. As a result, iMethyl had better predictive power than not only tumor mutation burden and PD-L1 expression but also genomic hypomethylation estimated by the array readouts. iMethyl particularly solves the issue of missing CpG methylation signals from non-CpG array probes.

However, tissue data suffers from low tumor purity. For example, low tumor purity causes inaccurate TMB estimates (22). DNA methylation measurements are also compromised by non-malignant cells such as adjacent normal and infiltrating immune cells resident in tumor tissues. In fact, DNA methylation is often used to assess the degree of tumor purity based on the extent to which genomic hypomethylation in tumor cells is diluted by non-malignant cells with normal methylation status (20,21,23–26). Such cellular heterogeneity should be accounted for by adjusting cell composition (27–30) to remove artefactual intertumoral variations in methylation.

Due to its minimal invasiveness and sample acquirement feasibility, cfDNA-based cancer screening is rapidly replacing tumor biopsies in the clinical setting. However, its application for cancer immunotherapy is in its early stages, despite cfDNA load (8–11), TMB (11,12), and copy number instability (14–16) having been identified as the predictors of ICB responses. In this work, we utilized the properties of genomic methylation in cfDNA for the first time in the field of cancer immunotherapy. Our assay based on ultra-deep sequencing allowed for the accurate estimation of methylation levels from cfDNA. We show that tumor purity problems can be overcome by measuring LINE-1 methylation from cfDNA. Furthermore, we demonstrate the feasibility of our method as a tool for evaluating ICB responses at early time points and monitoring disease progression during treatment. The robustness, accuracy, and reliability of our method as well as the previous predictors based on cfDNA should be validated in larger cohorts to be practically used in the clinical setting. Nonetheless, we hope that our method paves the way for reliable noninvasive prediction, early evaluation, and monitoring of clinical responses to ICB therapy.

## Materials and Methods

### Lung cancer ICB cohort

Advanced non-small cell lung carcinoma patients who were treated with anti-PD-1/PD-L1 monotherapy at Samsung Medical Center, Seoul, Republic of Korea were enrolled for this study. The present study has been reviewed and approved by the Institutional Review Board (IRB) of the Samsung Medical Center (IRB no. 2013-10-112, 2018-03-130, and 2018-04-048), and was conducted in accordance with the principles of the Declaration of Helsinki. All subjects provided their written informed consent, and there were no participants below 16 years. The following medical information was obtained: age, sex, stage, Eastern Cooperative Oncology Group (ECOG) performance status, pathology, comorbidity, smoking, treatment regimen, clinical response, and survival data. Lung tumor tissues were collected from the enrolled patients through bronchoscopy with or without endobronchial ultrasound or percutaneous needle biopsy. The tissues were then snap-frozen for storage at −80°C until use or stored as Formalin-Fixed Paraffin-Embedded (FFPE) blocks. Peripheral blood at the baseline was collected into commercially available EDTA-treated tubes. Plasma was separated from the entire blood by a density gradient centrifugation using the Ficoll-Paque™ PLUS (GE healthcare, Chicago, IL, USA), and plasma aliquots were stored at −80°C.

### Breast cancer ICB cohort

The KORNELIA trial was a multicenter, parallel-design, open-label phase 2 trial conducted in 10 academic hospitals in Republic of Korea (31). Ninety breast cancer samples were obtained from patients who were treated with nivolumab (anti-PD-1 antibody) until disease progression or unacceptable toxicity. The study has been reviewed and approved by the IRB of participating institutions, including Seoul National University Bundang Hospital (IRB no. B-1811-505-004), and was conducted in accordance with the principles of the Declaration of Helsinki. All subjects provided their written informed consent, and there were no participants below 16 years. The following medical information was obtained: age, sex, stage, ECOG performance status, pathology, comorbidity, smoking, treatment regimen, clinical response, and survival data. Baseline tumor biopsy from metastatic or recurrent lesions was required, and archival tumor samples were taken within 24 months before enrollment was allowed. Peripheral blood samples were collected at the baseline and at the time of progressive disease. The PD-L1 expression status was evaluated using SP263 antibody (Ventana Medical Systems) and scored as positive if tumor infiltrating immune cells were more than 1% stained. Peripheral blood at the baseline was collected into commercially available EDTA-treated tubes. Plasma was separated from the entire blood by a density gradient centrifugation using the Ficoll-Paque™ PLUS (GE healthcare, Chicago, IL, USA), and plasma aliquots were stored at −80°C.

### Ovarian cancer ICB cohort

A total of 74 patients with gynecological cancer who received immunotherapy at Yonsei Cancer Center, Seoul, Republic of Korea from December 2018 to January 2022 were enrolled in this study. Specifically, the cohort includes patients with ovarian cancer who received pembrolizumab or nivolumab monotherapy, or who received durvalumab with or without tremelimumab. Additionally, patients with cervical cancer who received tislelizumab or pembrolizumab were included. The study was approved by the institutional review board of Severance Hospital (IRB no. #4-2018-0342, #4-2018-0928). All subjects provided written informed consent, and there were no participants below 16 years. Clinical information including treatment regimen, duration of therapy, clinical response, and survival data was obtained.

### Evaluation of clinical response

The clinical response was evaluated by the Response Evaluation Criteria in Solid Tumors (RECIST v1.1) (32). The response to immunotherapy was classified into durable clinical benefit (DCB / responder) or non-durable benefit (NDB / non-responder). Complete response (CR), partial response (PR), or stable disease (SD) that lasted more than 6 months was considered as a DCB / responder. Progressive disease (PD) or SD that lasted less than 6 months was considered as an NDB / non-responder. Progression-free survival was calculated from the start date of treatment to the date of progression or death. Patients were censored at the date of the last follow-up for progression-free survival if they were alive without progression.

### iMethyl assay based on amplicon sequencing

Cell-free DNA was extracted from plasma using the QIAamp MinElute ccfDNA Kit (Qiagen) according to the manufacturer’s instructions. The extracted DNA was treated with bisulfite using the EZ DNA Methylation-Gold Kit (Zymo Research) according to the manufacturer’s instruction. Multiplexed PCR was performed by designing primers to target the 10 LINE-1probes. The sequences of the primer sets are provided in Supplementary Table 2. For multiplexed PCR, the primers for the target probes were divided into two groups: P3, P5, P8, P9, and P10 in one group, and P1, P2, P4, P6, and P7 in the other group. Multiplex PCR was performed for each probe group using the EpiTect MethyLight PCR Kit (Qiagen) in the following conditions: 5 min at 95°C followed by 23-29 cycles of 95°C for 15 sec and 60°C for 2 min. The 5’ ends of all PCR primers were phosphorylated. The PCR products in the two groups were mixed and were purified using the Agencourt AMPure XP PCR purification system (Beckman Coulter).

For the preparation of sequencing libraries, adaptors were ligated to the purified amplicons using the Quick Ligation Kit (NEB), and the ligation products were purified using the Agencourt AMPure XP system (Beckman Coulter). Finally, the library was amplified using the KAPA HiFi HotStart polymerase (KAPA Biosystems) with Illumina’s P5 and P7 sequencing primers in the following conditions: 2 min at 98°C followed by 7 cycles of 98°C for 15 sec and 64°C for 1 min. The amplified libraries were purified with the Agencourt AMPure XP system (Beckman Coulter), quantified with the KAPA Library Quantification Kit (KAPA Biosystems), and prepared for sequencing according to the standard normalization method described in “NextSeq 500 and NExtSeq 550 Sequencing Systems - Denature and Dilute Libraries Guide”. Paired-end sequencing of 150 cycles was performed using the NextSeq 550/550 Reagent Kit v2.5.

### Whole-exome sequencing

In this study, we conducted whole-exome sequencing for breast cancer and ovarian cancer, while the raw exome sequencing data for lung cancer was retrieved from Kim et al.’s study (33). Tumor samples were obtained before ICB treatment, and were then embedded in paraffin after formalin fixation or kept fresh. DNA was prepared using the AllPrep DNA/RNA Mini Kit (Qiagen, 80204), AllPrep DNA/RNA Micro Kit (Qiagen, 80284), or QIAamp DNA FFPE Tissue Kit (Qiagen, 56404) for library preparation for whole exome sequencing. Library preparation was performed by using SureSelectXT Human All Exon V5 (Agilent, 5190–6209) according to the instructions. Briefly, 200–300 ng of tumor and normal genomic DNA was sheared, and 150–200 bp of the sheared DNA fragments were further processed for end-repairing, phosphorylation, and ligation to adaptors. Ligated DNA was hybridized using whole-exome baits from SureSelectXT Human All Exon V5. The libraries were quantified by Qubit and 2200 Tapestation, and sequenced on an Illumina HiSeq 2500 platform with 2 × 100 bp paired ends. The target coverage for the normal samples and tumor samples was × 50 and x 100, respectively. The sequencing data was aligned to the hg19 reference genome using the Burrows-Wheeler Aligner mem module (v.0.7.17) (34). The data was further filtered by marking and removing duplicate reads using Picard (http://broadinstitute.github.io/picard, v.2.26.10). The base quality score was recalibrated with the Genome Analysis Toolkit (GATK v.4.2.4.1) (35). Tumor mutation burden (TMB) was calculated as the number of amino acid-changing somatic mutations called with MuTect2 with a matching panel of normal. To find neoantigen candidates, the mutations were annotated with the Ensembl Variant Effect Predictor (VEP), and the amino acid sequence neighboring the mutation was performed with pVACseq (v.4.0.10) (36). The HLA typing was conducted with OptiType (37), and the prediction of MHC binding neoantigens was conducted with NetMHCpan (v.4.1) (38). Neoantigen load (NeoAg) was obtained as the total number of the identified neoantigen candidates. Tumor purity was estimated with the exome data by using Sequenza (v3.0.0) (39) and ABSOLUTE (20).

### Methylation array profiling

The methylation array data was obtained from our previous work conducted by Jung et al. (17). Array-based tissue methylation profiling was performed by following the instructions of the Infinium MethylationEPIC BeadChIP Kit (Illumina, WG-317-1002). Briefly, 500 ng genomic DNA (gDNA) was used for bisulfite conversion using the EZ DNA methylation kit (Zymo Research, D5001). The bisulfited gDNA was denatured and neutralized for amplification, and was further processed for fragmentation. After fragmentation, DNA was eluted and resuspended in a hybridization buffer, and then hybridized onto the BeadChip. The BeadChip was prepared for staining and extension after washing out the unhybridized DNA, and it was imaged using the Illumina iScan System. The raw intensity files were then preprocessed into beta values using the preprocessIllumina function in minfi (40). The PMD levels of our cohort samples were calculated based on the average of the EPIC probes for Solo-WCGW CpGs in common PMDs (18) (provided at https://zwdzwd.github.io/pmd). Redundant probes such as multi-hit probes were filtered by using the filter function of the ChAMP package (41).

### iMethyl data processing

More than 650,000 raw read pairs per sample (∼100 Mb raw output) were generated with a NextSeq550Dx machine in the 2×75 bp mode (150 cycles). Raw sequence reads were adaptor-trimmed with cutadapt (42) (v2.8, --minimum-length 30). Based on the human reference genome hg19, bisulfite-converted (C:G>T:A) template sequences were prepared with bismark_genome_preparation (bismark v0.22.3) (43). Preprocessed reads were aligned into bisulfite-converted genomes by using bismark (43) with bowtie2 (44) (v2.3.5) as a genome aligner with the default options. Aligned reads were sorted using samtools (v1.9) (34,45) and the reads with indels near target regions +/− 30 nt were removed. After the filtering and processing, the average read count per sample was 499,533 (lung cancer iMethyl-liquid, n=167), 658,009 (lung cancer iMethyl-tissue, n=137), 436,315 (breast cancer iMethyl-liquid, n=91), 486,209 (breast cancer iMethyl-tissue, n=50), and 930,036 (ovarian cancer iMethyl- liquid, n=74).

The following steps were followed to calculate the beta value for each iMethyl probe. The reads aligned to the Watson- or Crick-strand were separated, and then a C-to-T transition from the Watson-strand and G-to-A transition from the Crick-strand was counted for each target probe using bam-readcount (46) (v0.8.0) and our in-house script. The ratio of the methylated (C or G) base at each probe was calculated and corrected with the sample conversion rate.

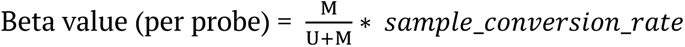

M: number of methylated bases at the target probe (C for Watson, G for Crick)

U: number of methylated bases at the target probe (T for Watson, A for Crick)

To estimate the sample conversion rate, or the rate of base conversion by bisulfite treatment at the level of samples, 92 naïve cytosine (i.e., non-CpG site) positions across the probe sequences were utilized. The rates of bisulfite conversion (C-to-T or G-to-A) at the 92 positions were averaged to obtain the sample conversion rate. The average sample conversion rate in each cohort was 0.998 (lung cancer iMethyl-liquid), 0.995 (lung cancer iMethyl-tissue), 0.993 (breast cancer iMethyl-liquid), 0.993 (breast cancer iMethyl-tissue), and 0.989 (ovarian cancer iMethyl-tissue).

Among the 137 lung cancer samples subjected to iMethyl-tissue, 8 samples were missing beta values for P8, P9, and P10. To compensate for the missing values, we performed imputation. With the 129 samples whose methylation values were complete, we built three multiple linear regression models with each of the missing probes as the response variable and the 7 probes (P1-P7) as the explanatory variable. Using the models, the missing beta values were predicted by fitting them against the 7 probes (P1-P7). This analysis was performed using the lm function of the R package.

### Cohort survival analysis

To test the power of iMethyl as an ICB response prediction marker, we performed the survival analysis in different conditions. The methylation-high and -low group was defined according to the mean value of each cohort. For a more robust analysis, we implemented bootstrapping by generating 1,000 or 5,000 resamples and conducting the survival analysis using each resample. If the methylation-high group showed better survival, which coincided with our hypothesis, the P value was used as-is. On the contrary, if the methylation-low group showed better survival, we used the negative P value to ensure that both cases were considered. Depending on data availability, the lung and breast cancer cohorts used overall survival data whereas the ovarian cancer cohort used progression free survival data. The survival analysis was performed based on the Kaplan-Meier curve using the survival (v3.1.12) (45) and survminer (v0.4.9) (43) packages in R. The survival graphs were illustrated with the ggplot2 R package (v3.3.5) (47). The survival analysis was performed based on the Kaplan-Meier curve, and the difference between the methyl-high and -low group was analyzed by the log-rank test. Wilcoxon signed-rank test was used to compare the statistical powers of predicting clinical outcomes of ICB therapy for iMethyl. P values less than 0.05 were considered significant.

### Probe weighting

Probe weights were computed to maximize the predictive power of iMethyl for ICB responses. To obtain optimal weights, we generated simulated probe sets that consist of randomly chosen iMethyl probes. We prepared 10 probe sets of different sizes (i.e., 10, 15, 20, 30, 50, 75, 100, 150, 200, and 300). The weights were assigned as the number of each probe in each simulated set divided by the size of the set. For example, if P1 was chosen 6 times in the simulated set size of 30, the weight assigned to P1 would be 0.2. For each simulated set, random sampling was conducted for 5,000 iterations to achieve statistical robustness. Finally, the resulting 50,000 sets of weights (5,000 iterations for each of the 10 simulated sets) were applied to the corresponding beta values for the stratification of patient samples according to the average methylation level in the survival analysis. The set of weights with the best performance was selected.

### Principal component analysis and hierarchical clustering

For the principal component analysis, we calculated eigenvectors and eigenvalues of the 10 probes using singular value decomposition in the scaled beta value matrix using the prcomp R function (50). Then, the PCA plot for the two first principal components with the largest variance was plotted with the factoextra R package (51). The hierarchical clustering was performed across the samples and probes, and the heatmaps were plotted using the pheatmap R package (v1.0.12) (52).

## Author’s contributions

K.H.K., H.M.K., I.K.S. and S.-J.N. were equally contributed to this work. K.H.K. performed all data analyses and wrote the manuscript with J.Y.K. H.M.K., K.J.S., Y.-N.K. and J.-Y.L. generated and managed the cohort data. I.K.S., S.-J.N. and D.-Y.C. designed the iMethyl assay and calculated the beta value for each iMethyl probe. S.H.K, J.H.K and S.-H.L. supervised the cohort analysis. J.K.C conceived and supervised the whole study.

## Data availability

All iMethyl data produced in this work is provided in the Supplementary Tables. For lung cancer ICB cohort, Jung et al.’s methylation chip data was available at Gene Expression Omnibus under GSE119144 and Kim et al.’s raw exome sequencing data was retrieved with the accession number EGAS00001002556. The raw exome sequencing data of our other ICB cohorts have been submitted to the European Genome-phenome Archive (EGA) under accession number EGAS00001007490 (https://wwwdev.ebi.ac.uk/ega/studies/EGAS00001007490) for breast cancer and EGAS00001007489 (https://wwwdev.ebi.ac.uk/ega/studies/EGAS00001007489) for ovarian cancer.

## Competing Interests

The authors declare that they have no competing interests.

## Funding

This research was supported by the Bio & Medical Technology Development Program of the National Research Foundation of Korea funded by the Ministry of Science and ICT (NRF- 2017M3A9A7050612 and NRF-2019M3A9B6064688). This work was also supported by grant no. 16-2020-0004 from the SNUBH Research Fund.

## Notes

### Competing Interest Statement

The authors have declared no competing interest.

