## Supplementary material for "Genomic hypomethylation in cell-free DNA predicts responses to checkpoint blockade in lung and breast cancer": Figures S1-S12

### Supplementary Figure 1

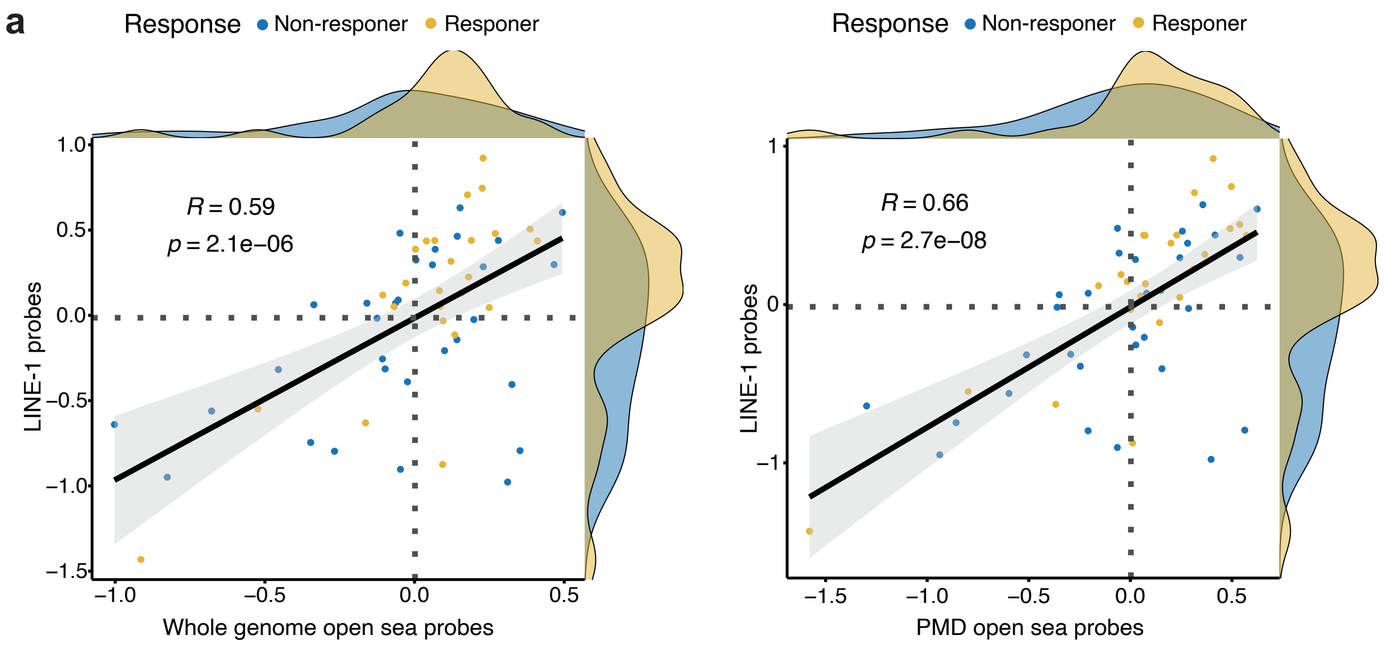


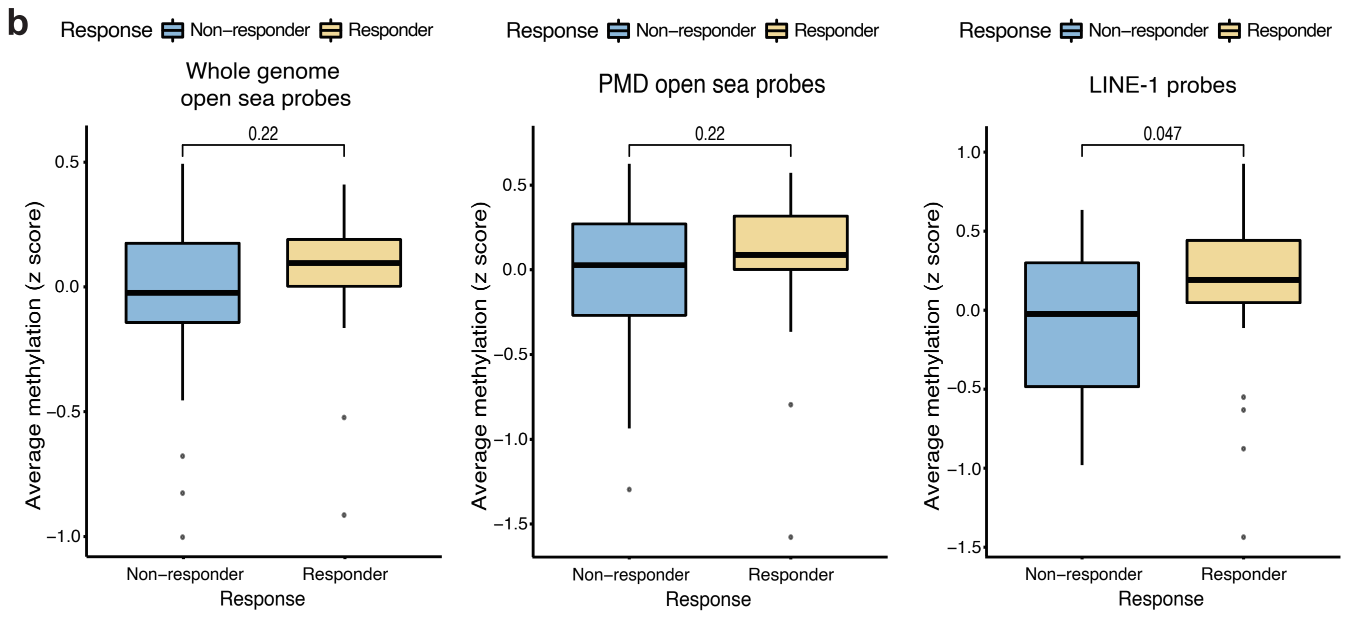

#### Supplementary Figure 2

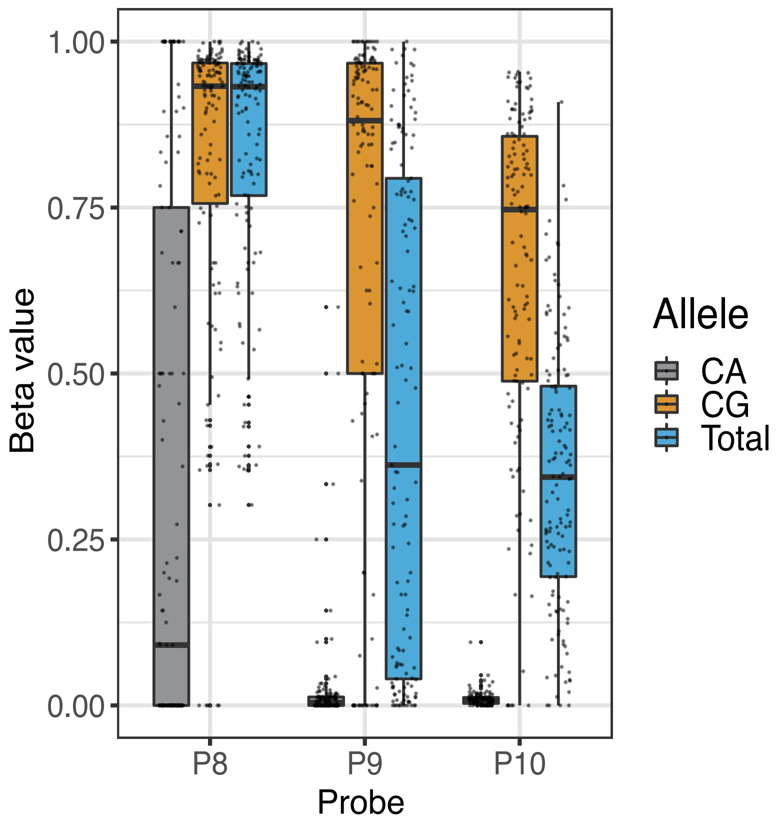

### Supplementary Figure 3

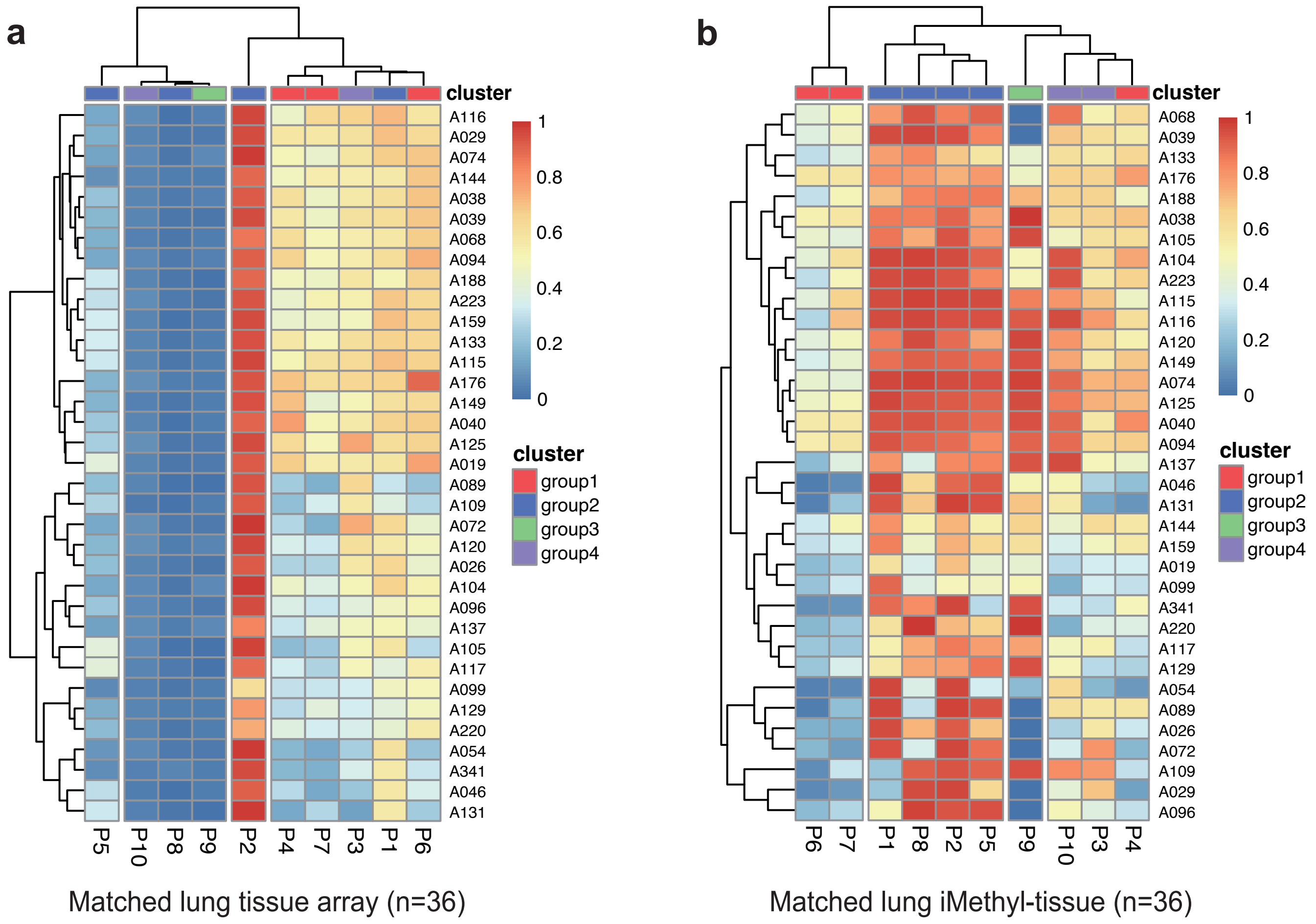

### Supplementary Figure 4

a

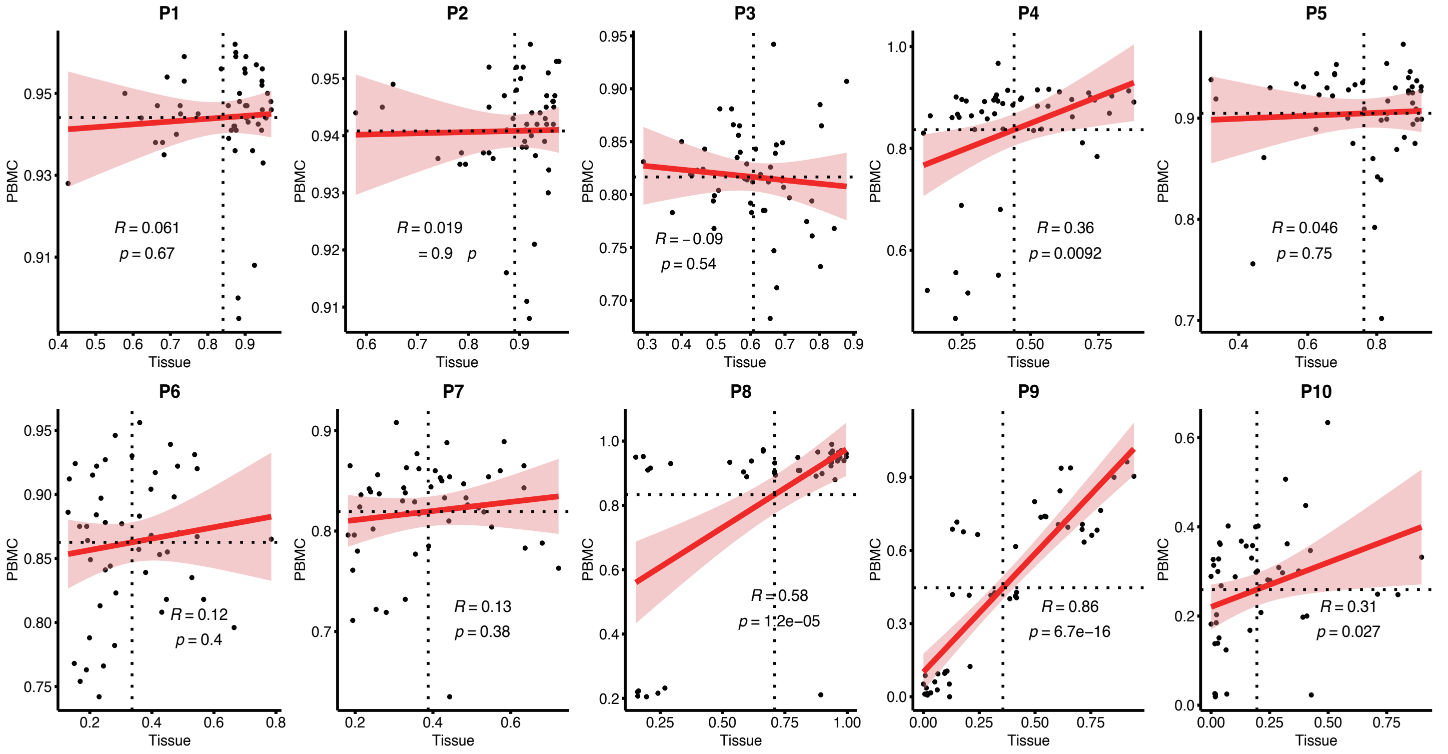


b

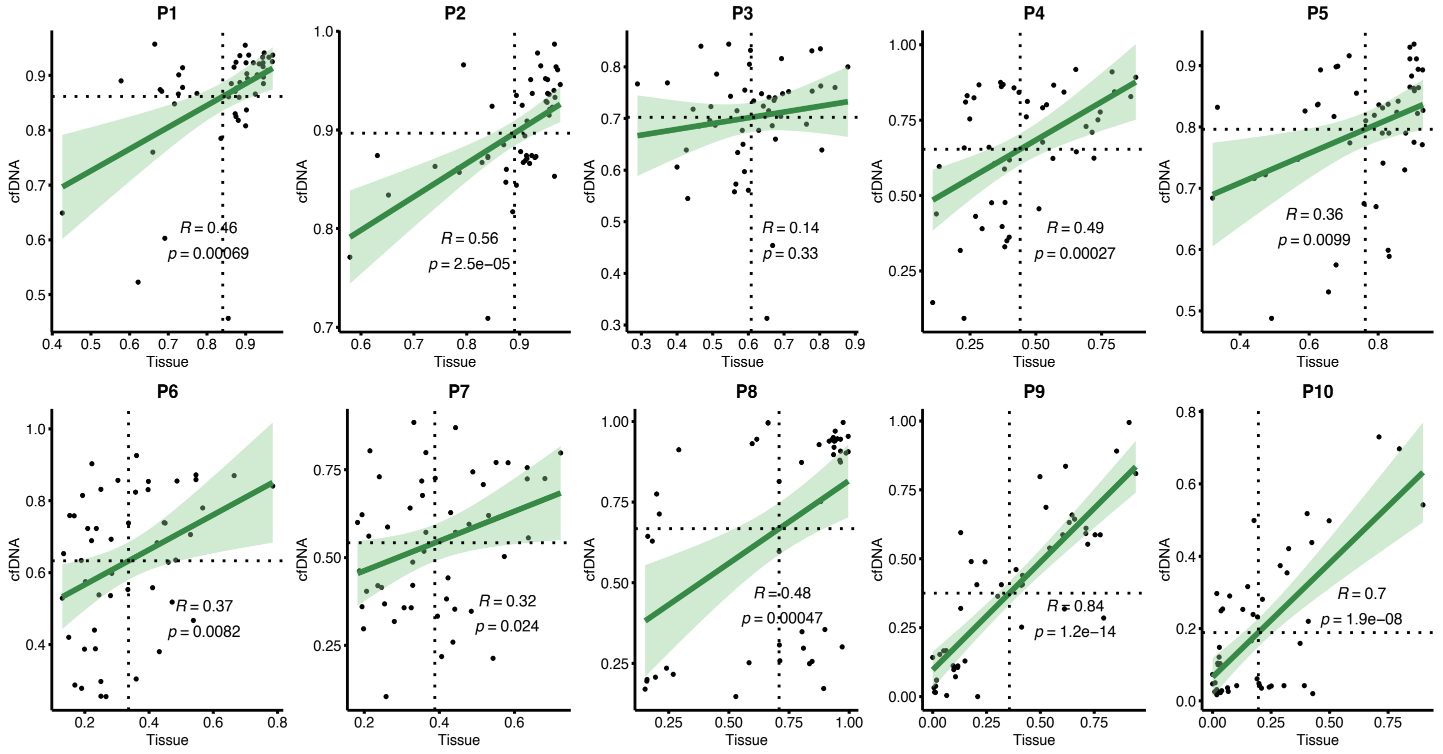

### Supplementary Figure 5

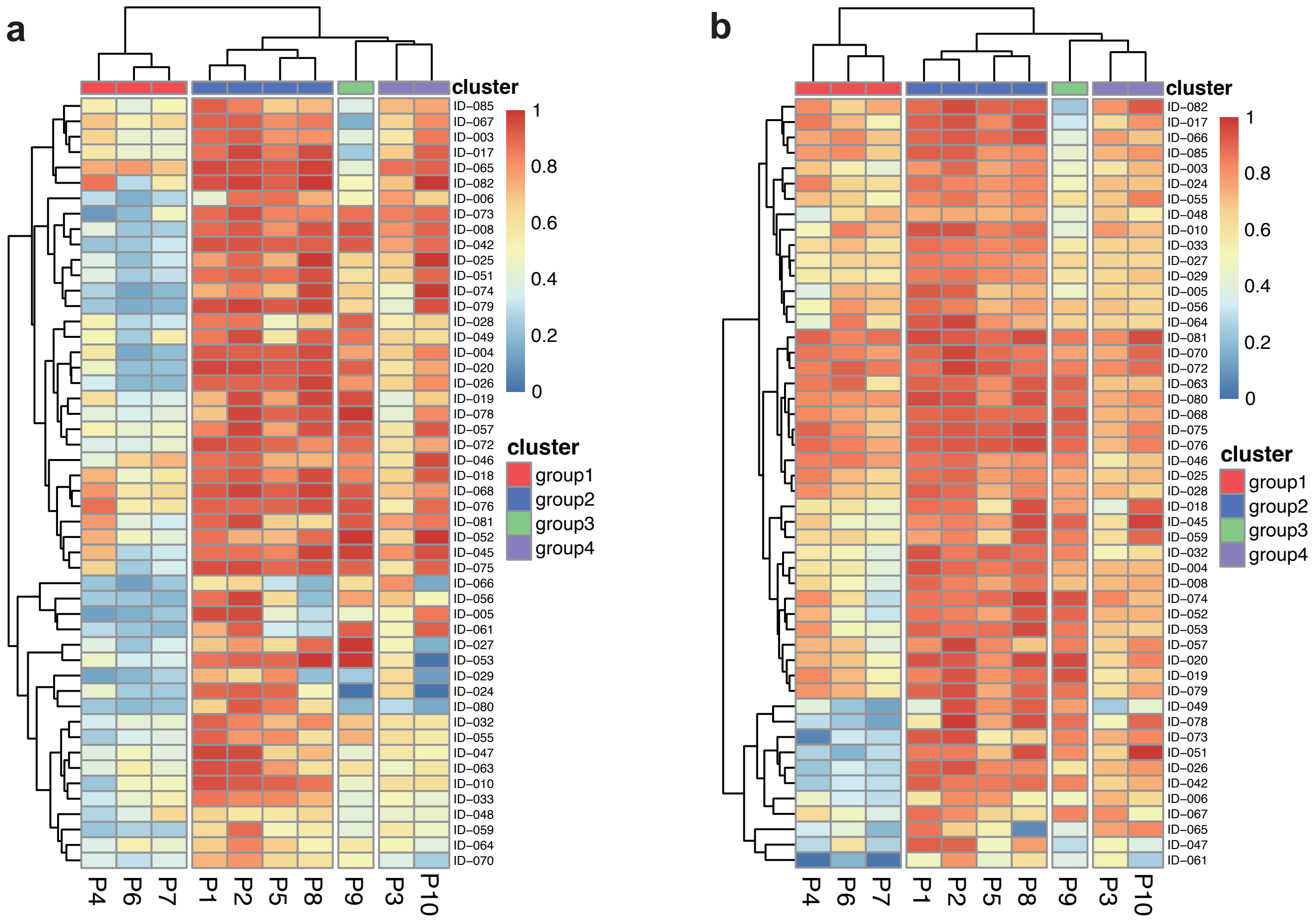


Matched breast iMethyl-tissue (n=50)

Matched breast iMethyl-liquid (n=50)

**Supplementary Figure 6**

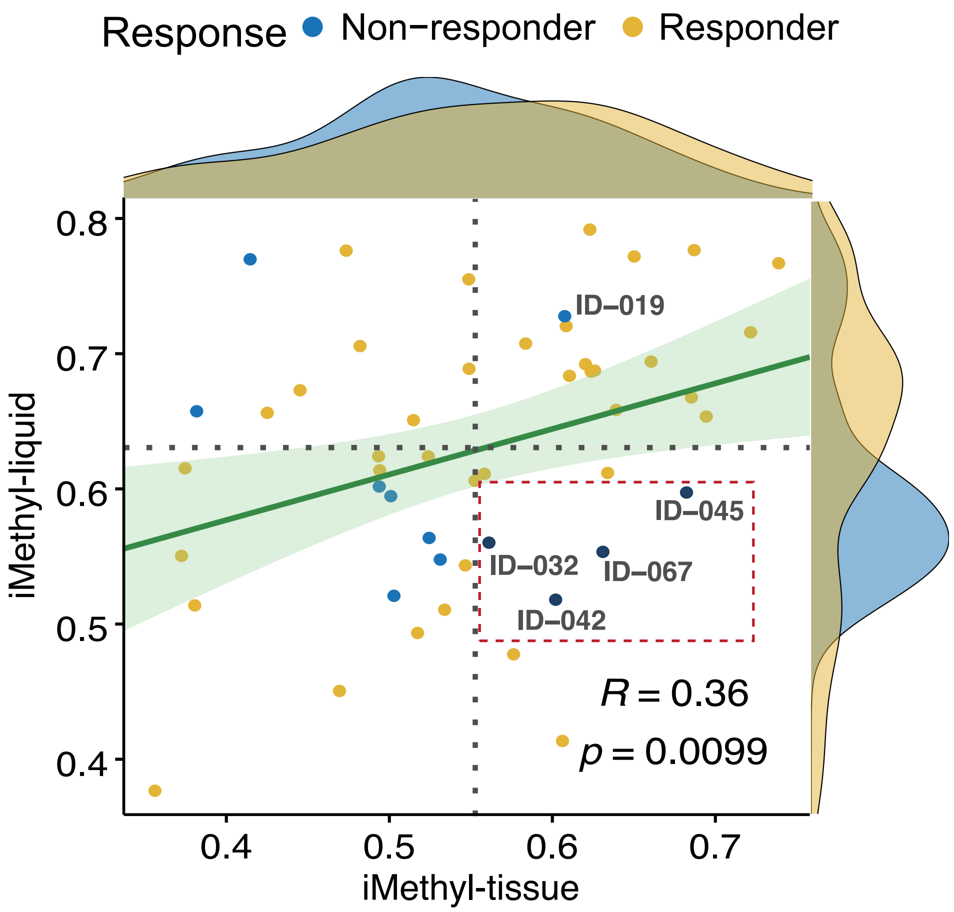

### Supplementary Figure 7

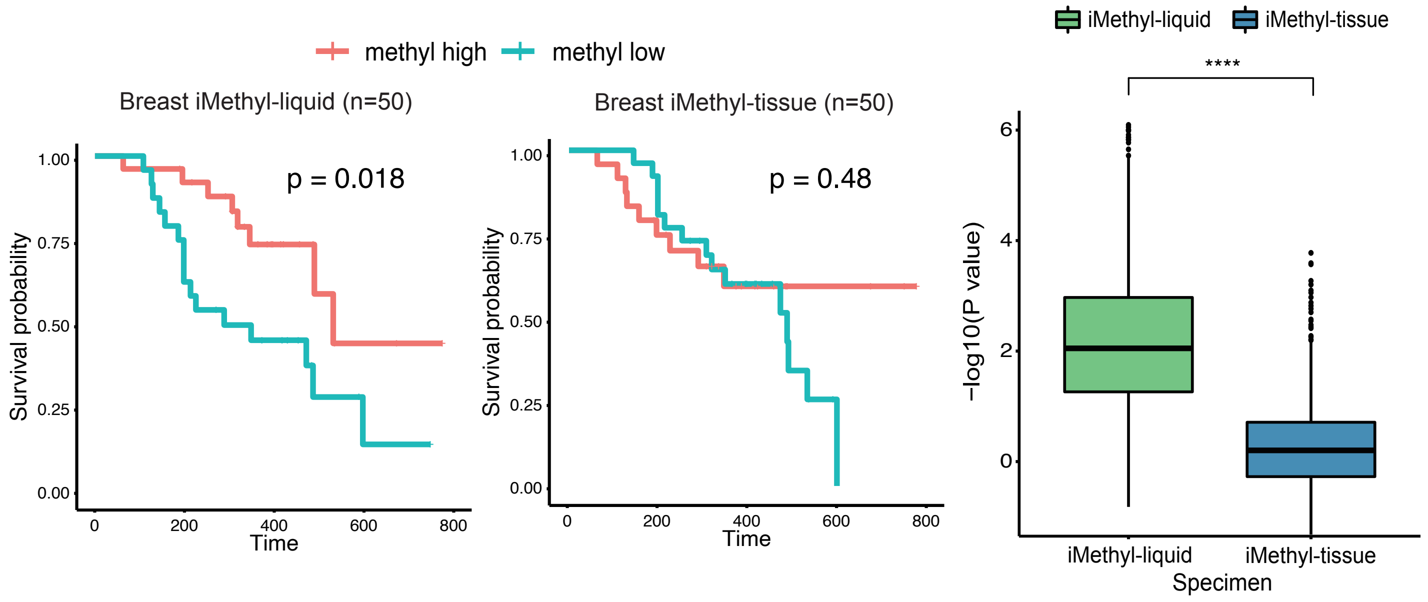

### Supplementary Figure 8

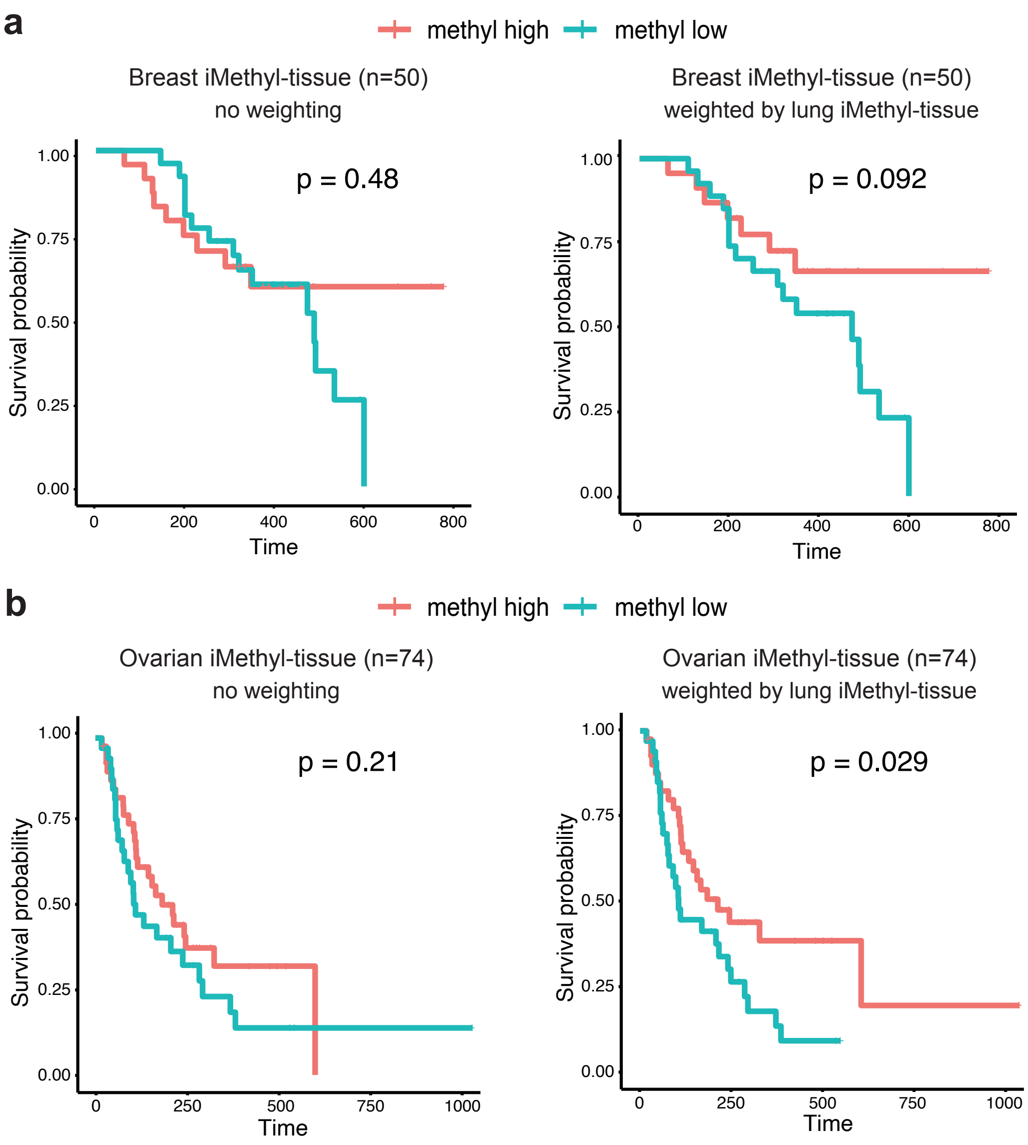

### Supplementary Figure 9

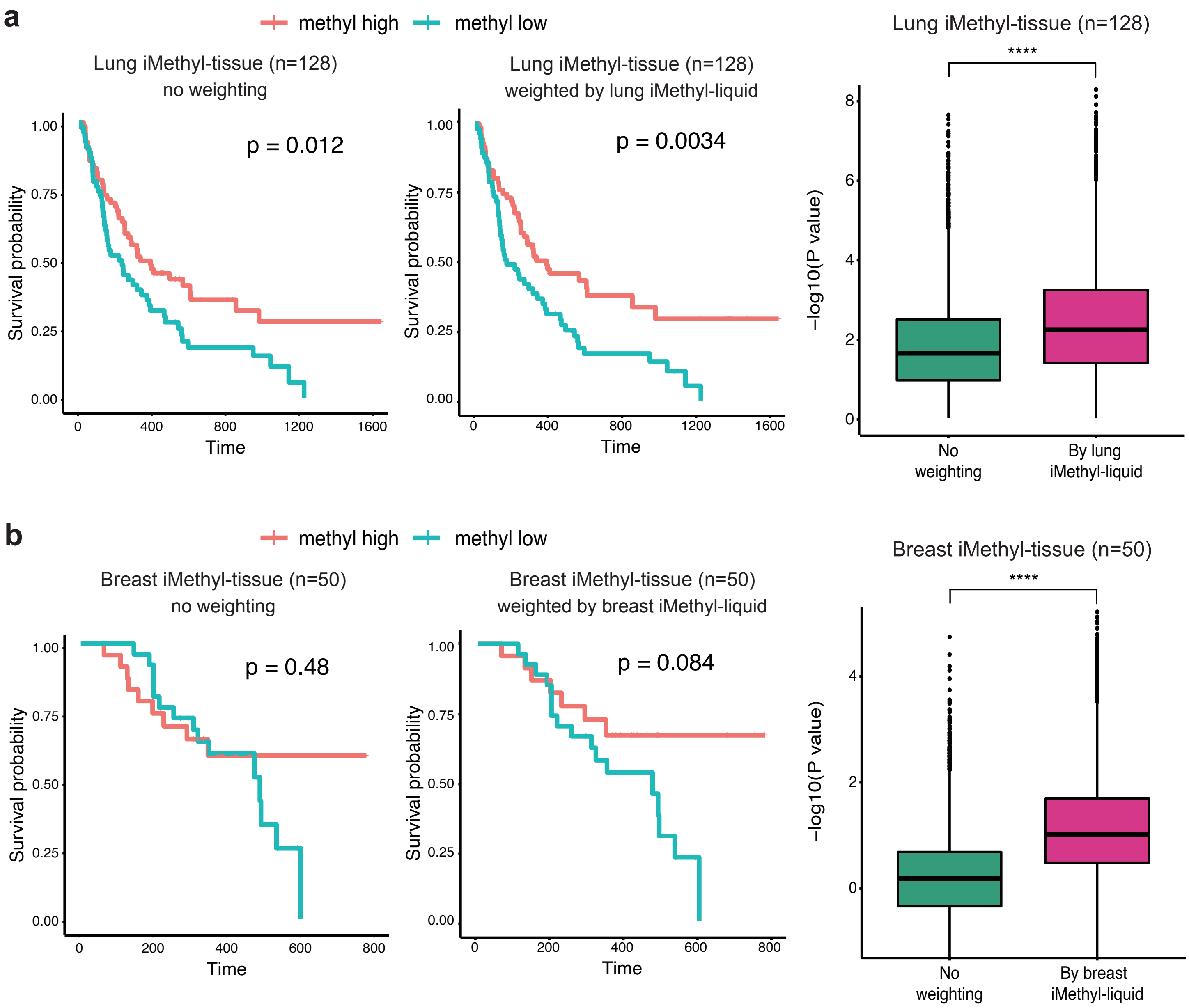

### Supplementary Figure 10

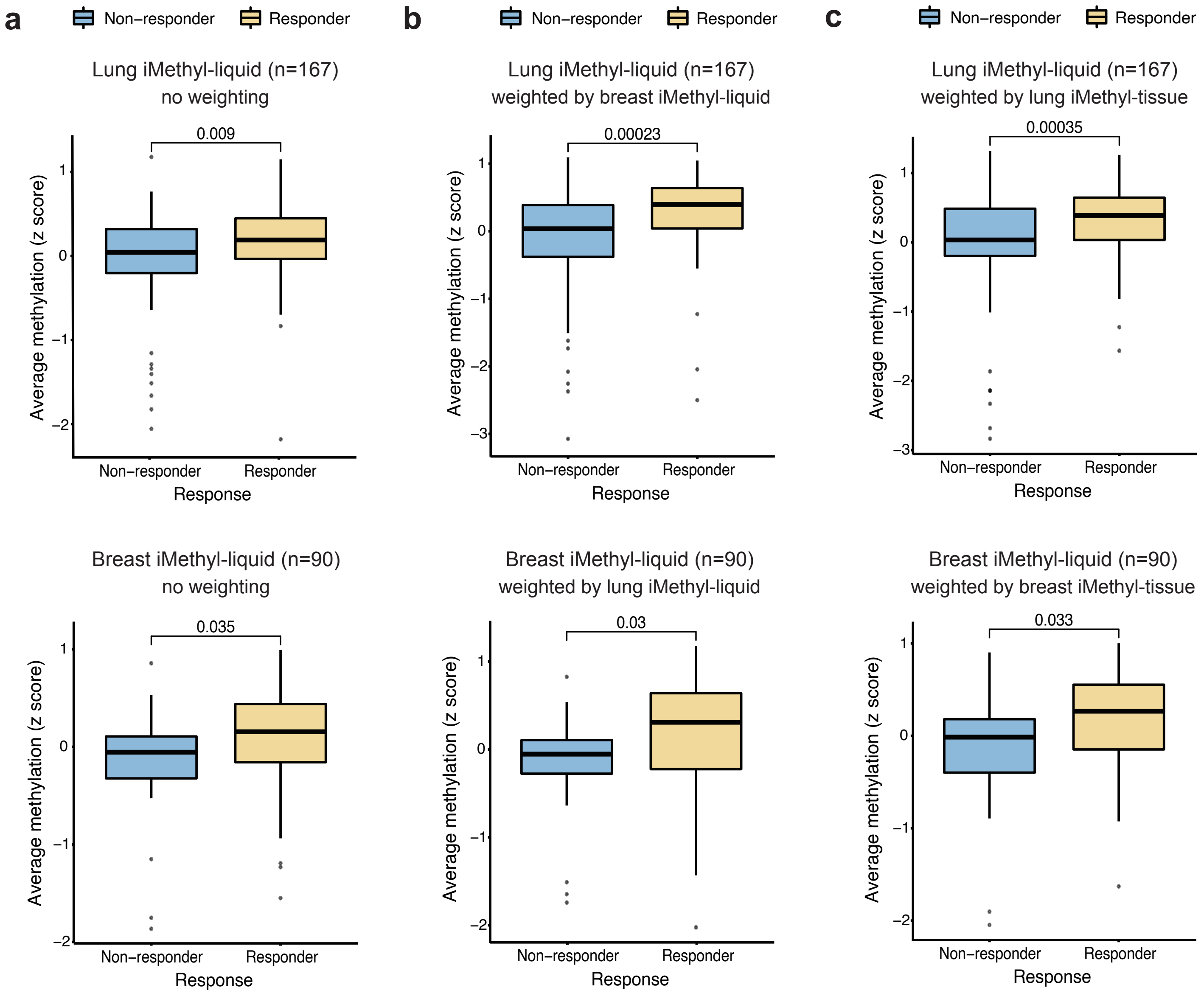

### Supplementary Figure 11

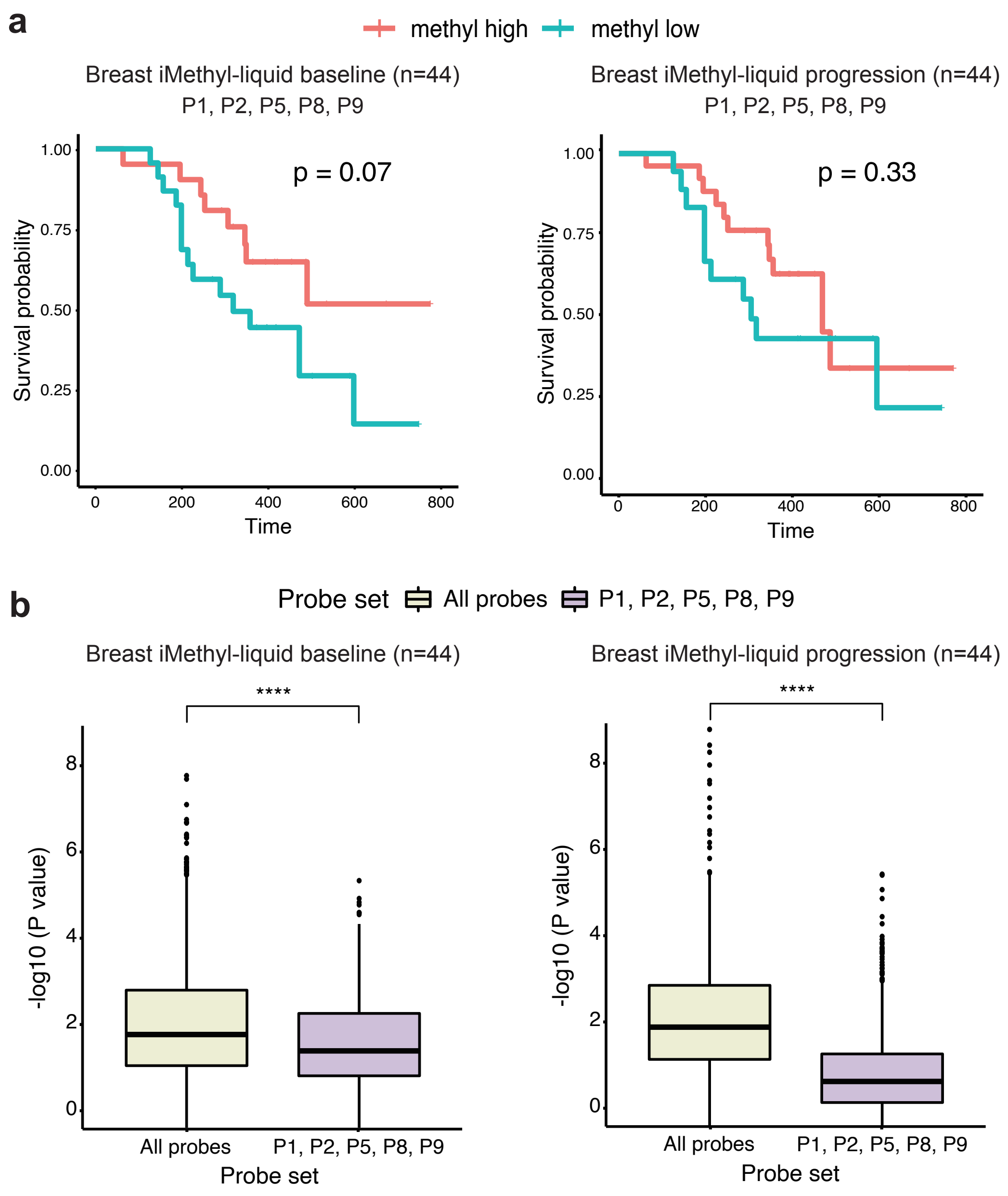

### Supplementary Figure 12

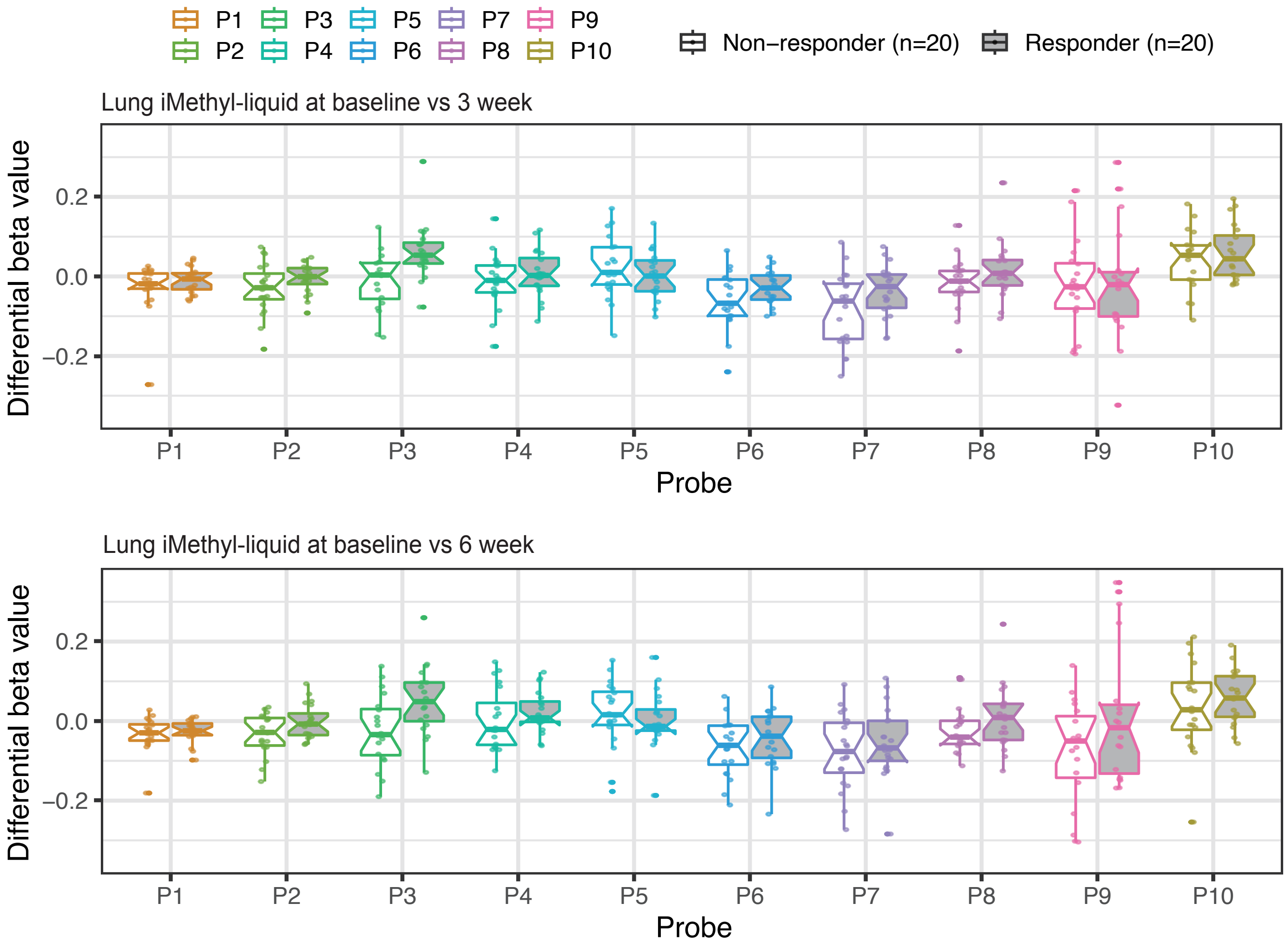
